# An in vivo platform to jointly monitor cellular and metabolic responses to chemotherapy

**DOI:** 10.64898/2026.06.25.734296

**Authors:** Veronika Pister, Zuzana Tatarova, Noel Park, Gautami Gaidhani, Juraj Jakubik, Laura Heiser, Jenna Blum, Alec Palmiotti, Ellen Maloney, Ernest Fraenkel, Shawn M Davidson, Oliver Jonas

**Affiliations:** Department of Biological Engineering, Massachusetts Institute of Technology, Cambridge, MA 02139, USA; Georg-Speyer-Haus Institute for Tumor Biology and Experimental Therapy, Frankfurt, Germany; German Cancer Research Center and German Cancer Consortium, Heidelberg, Germany; Frankfurt Cancer Institute, Goethe University Frankfurt, Frankfurt, Germany; Department of Molecular Biology, Princeton University, Princeton, NJ, USA; Department of Biomedical Engineering, Oregon Health and Science University, Portland, Oregon; Knight Cancer Institute, Oregon Health and Science University, Portland, Oregon; Department of Medicine, Division of Pulmonary and Critical Care Medicine, Feinberg School of Medicine, Northwestern University, Chicago, IL 60611, USA; Northwestern University Interdepartmental Neuroscience (NUIN), Northwestern University, Chicago, IL 60611, USA; Department of Radiology, Brigham and Women’s Hospital, Harvard Medical School, 221 Longwood Ave, Boston, MA 02115, USA; Broad Institute of MIT and Harvard, Cambridge, MA 02139, USA; Robert H. Lurie Comprehensive Cancer Center, Northwestern University, Chicago, IL 60611, USA; Center for Human Immunobiology, Northwestern University, Chicago, IL 60611, USA

## Abstract

How drug treatments reshape immune and metabolic states within intact tumors remains difficult to study with existing methods. We introduce a spatial pharmacology platform that enables parallel analysis of multiple agents within a single tumor, linking local drug exposure to immune and metabolic remodeling. Using a microdevice for localized drug delivery, we created a large-scale paired CyCIF–MALDI dataset spanning 1.5 million cells across 27 MMTV-PyMT tumor sections and nine treatment programs, enabling integrated spatial pharmacology at unprecedented scale. Metabolic signatures robustly predict proteomic spatial neighborhoods establishing metabolism as a powerful predictor of tumor organization and immune phenotype. Within this framework, we identify a dominant metabolic axis defined by the myeloid polarization between CSF1R+ tumor-associated macrophages and MPO+ infiltrating myeloid cells localized near regions of drug-induced tumor cell death. Finally, we detect putative lipid-associated macrophage (LAM)-like populations within drug-resistant treatment regions.

## Main

Metabolic conditioning of the tumor microenvironment (TME) is a central determinant of immune cell function and contributes to therapeutic resistance^1^. Immune cells are disrupted by the metabolic TME through two mechanistically distinct axes: direct fate-altering signals that repolarize innate immune cells toward pro-tumor states, and nutrient competition that drives functional exhaustion. Tumors accumulate immunomodulatory metabolites that suppress antigen presentation and cytotoxic effector programs via acidosis-driven lactate signaling^2,3^, adenosine–A2A receptor signaling^4,5^, and the kynurenine–AHR axis^6–8^. In parallel, nutrient competition between tumor cells and immune cells constrains T-cell glycolysis and IFN-γ production in vivo, directly linking tumor bioenergetics to T-cell dysfunction^9–11^.

Resolving immunometabolic states within intact tissue remains challenging. Bulk metabolomic assays require large cell numbers and average signals across heterogeneous microanatomical niches. In addition, cell isolation and fluorescence-activated cell sorting perturb cellular metabolism and introduce metabolomic artifacts^12–14^. Finally, metabolic phenotypes observed in standard culture conditions often diverge from in vivo behavior, even when partially mitigated by physiologic media formulations^15,16^.

Recent efforts to overcome these limitations using in situ or in vivo spatial metabolite imaging have provided valuable insights but studies either lack the paired proteomic context necessary for immune cell-type distinction^17,18^, or are limited in spatial and molecular coverage to capture the diversity of immunomodulatory metabolism^19–21^. Critically, no prior study has resolved immune cell–type–specific metabolic remodeling across defined therapeutic perturbations within intact tumors.

To investigate immune phenotypes in regions of therapeutic response and resistance, we pair intratumoral microdelivery of therapeutics with high-dimensional MALDI-MSI to obtain spatially resolved readouts of tumor microenvironment remodeling adjacent to each drug reservoir. This multiplex implantable microdevice (IMD) assay (MIMA) resolves treatment-specific immune and stromal signatures and has previously predicted rational drug combinations in vivo^22,23^ and in patients^24–26^. We selected perturbations spanning distinct immune response and those with highest tumor killing capacity based on prior MIMA readouts. For example, Venetoclax recruits immature myeloid populations; Panobinostat promotes immunogenic cell death and increases antigen-presenting myeloid activity^22,27,28^; the combination (panobinostat/venetoclax, PV) shows the strongest in vivo efficacy (Xu *et al.*, under review).

We applied this in the MMTV-PyMT genetically engineered model of luminal-like breast cancer, which develops in an immunocompetent setting with macrophage-rich tumor microenvironments characteristic of human disease^29,30^. This enables spatially resolved, causally interpretable assessment of how pharmacologic perturbations reshape local immune ecologies under native metabolic constraints, positioning immunometabolism as a measurable and actionable determinant of therapeutic response.

Together, we establish an end-to-end spatial pharmacology pipeline for the systematic investigation of immunotherapeutic agents. By integrating intratumoral microdelivery with paired immunofluorescence and metabolomic imaging, cross-modal registration, and spatial colocalization analysis, this framework enables causal readouts of drug-induced immunometabolic remodeling within intact tumors.

## Results

### An implantable microdevice enables controlled, multiplexed drug perturbation in intact tumors

We deployed an implantable microdevice (IMD)^31,22,24,25^ bearing 18 spatially segregated release sites to perturb intratumoral regions within single tumors (Fig. 1a). Small-molecule agents were formulated in poly(ethylene glycol) (PEG) for precise intratumoral release and loaded to approximate tissue-level exposures corresponding to systemic dosing (Methods). PEG-only control wells and untreated intratumoral regions served as local controls. IMD placement and retrieval were well tolerated by animals bearing orthotopic breast tumors, with the architecture surrounding release sites preserved on hematoxylin and eosin (H&E) sections, allowing downstream multi-omic profiling without tissue dissociation (Fig. 1b).

**Figure 1:**
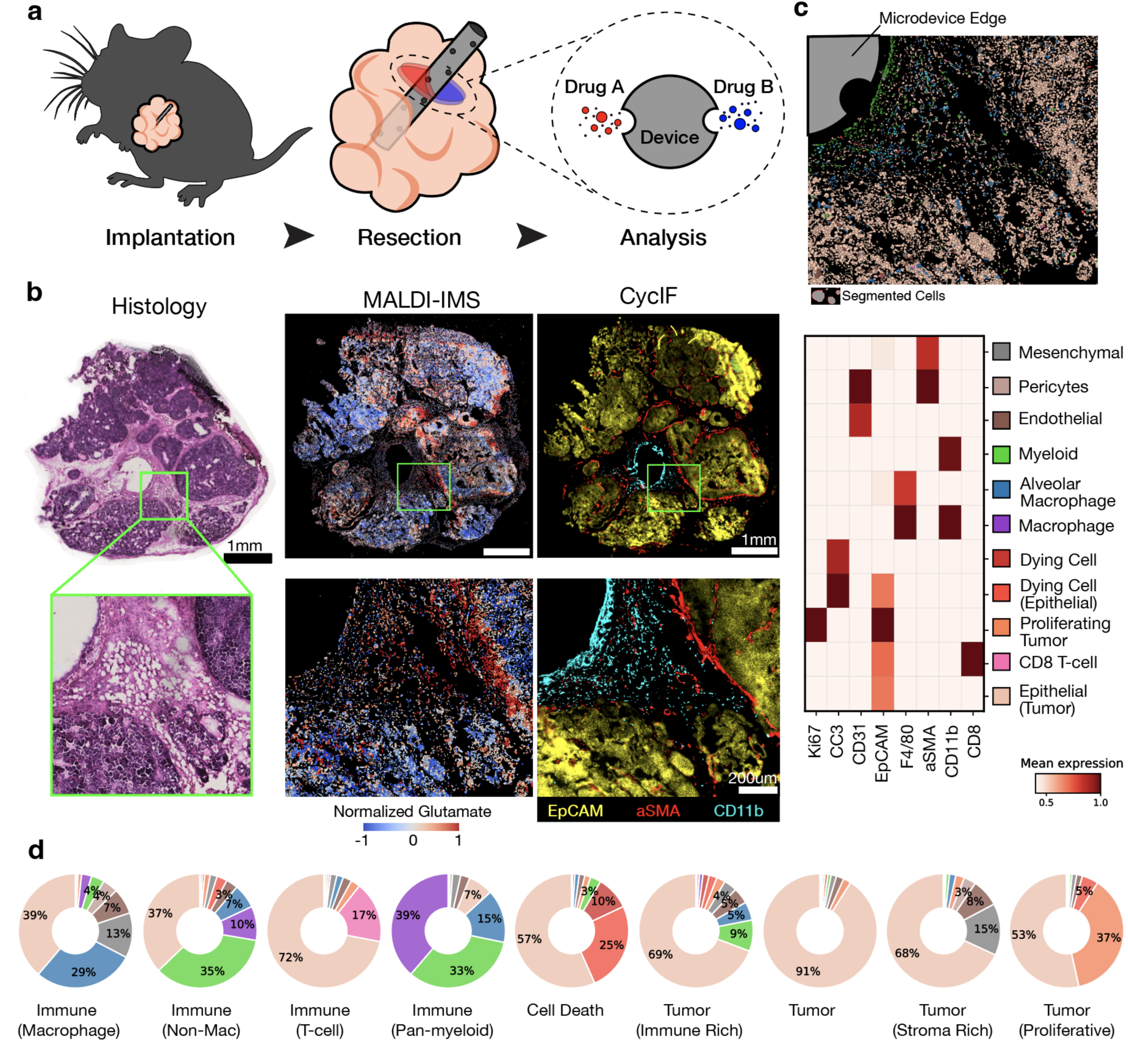
In vivo multi-omic profiling of drug-induced immune phenotypes using an implantable microdevice. (a) Schematic of the nanonail implantable microdevice (multiplex implantable microdevice assay, MIMA) configured with up to 18 spatially separated release sites, enabling parallel perturbation of discrete intratumoral regions within a single tumor. Each treatments’ associated loading location, mechanism of action, and the expected phenotypic response in adjacent tissue is displayed. (b) Overview of the multimodal paired dataset. For each experimental condition on the device, two serial sections yield 10 CycIF images, 84 metabolite images from MALDI-IMS, and an H&E section corresponding to the MALDI-analyzed tissue. The MALDI-IMS panel shows the grid of MALDI-IMS detection masked by the cell segmentation from the CycIF DAPI image. (c) Proteomic marker expression profiles across resolved cell types. Marker intensity (x-axis) is normalized column-wise to a maximum of 1, and cell types are shown along the y-axis. (d) Cellular neighborhoods derived from CycIF-based cell–cell spatial relationships. Nine neighborhoods were identified, comprising four immune neighborhoods, four tumor neighborhoods, and one cell-death neighborhood; the fractional cell-type composition of each neighborhood is shown. Legend as in c.

Following a three-day exposure interval, tissues adjacent to each release site were sectioned serially for cyclic immunofluorescence (CycIF)^22^ and matrix-assisted laser desorption/ionization imaging mass spectrometry (MALDI-IMS), with an intervening H&E reference (Fig. 1b). We registered the modalities by thin-plate spline (TPS)^32^ warping using conserved histologic features as landmarks, yielding a common coordinate framework for metabolites and single-cell phenotypes. TPS significantly outperformed alternative rigid and affine registration methods for both within-modality serial-section registration and cross-modality alignment of CyCIF-derived cell death regions with histologically labeled regions (paired t-test, P < 0.01; Extended Data Figs. 1, 2).

### CycIF identifies drug-recruited immune cellular neighborhoods in situ

We profiled 20 sections across eight treatment programs with a validated 8-antibody CycIF panel for cell type identification (Fig 1c, Extended Data Fig. 3a). 14 sections were additionally imaged with an expanded 9-antibody panel to refine cell-state annotation^22,33^. After preprocessing, nuclei were segmented and single-cell intensities quantified. Marker positivity was gated using Gaussian mixture-modeling to minimize operator bias while accommodating non-Gaussian tails (Methods). This strategy is consistent with model-based gating in cytometry and multiplexed imaging^34,35^. Adhering to canonical patterns of marker expression, we were able to label 1.48 million cells with cell type identities (Extended Data Fig. 3b).

Spatial neighborhood analysis summarizes recurrent patterns of local cell type organization^36–38^. Using this established approach (Extended Data Fig. 3c), we assigned each cell to a neighborhood based on the composition of its surrounding cells (Fig. 1d and Methods). We identified nine robust proteomic neighborhoods, including four that were immune-enriched (>20% immune cells): Immune (macrophage), Immune (non-macrophage), Immune (pan-myeloid), and Immune (CD8 T cell). Four non-immune neighborhoods were tumor-dominated (>70% tumor cells): Tumor, Tumor (stroma enriched), Tumor (immune-enriched), and Tumor (proliferative). Finally, one neighborhood was enriched for CC3+ cell death (35% dying cells): Cell Death (Fig. 1d, Extended Data Fig. 3d).

### MALDI-IMS resolves spatially structured metabolic programs across tumor compartments

To determine the metabolic profiles of different immune neighborhoods, we performed spatial metabolomic profiling on 27 MMTV-PyMT tumor sections using MALDI-IMS. After preprocessing, the dataset comprised 230,812 spots, which were normalized by total ion current (TIC), log-transformed, and z-scored across each metabolite (Methods). 84 high-quality small molecule m/z features were detected across all sections and retained for downstream analysis. Putative metabolite identities were assigned using high-precision matching of mass-to-charge ratios; orthogonal confirmation is provided where indicated in Methods (Extended Data Fig. 4). Six identified lipid signatures were labeled by empirical chemical formula. Isobaric species (e.g., citrate and isocitrate) are indistinguishable by MALDI-IMS and were therefore reported as a single peak.

Tissue sections were organized by coordinated shifts in groups of co-abundant metabolites. To quantify spatial coupling patterns between metabolites, we constructed a spatial correlation network (Methods). This network resolved three robust spatially correlated metabolic programs, defined by correlations in the top decile of Bivariate Moran I statistics that were mutually conserved across all mouse replicates: (i) glutamine-nucleotide metabolic coupling, (ii) lipid metabolism, and (iii) purine catabolism. In addition, spatial anticorrelations between palmitic acid and both taurine and glutamine were conserved across all replicates.

To interpret these metabolic programs, we examined metabolite abundance across tumor compartments (Extended Fig. 5a). A board-certified pathologist (SA) annotated the MALDI-IMS slide to delineate tumor, stroma, muscle, necrotic, and cystic regions. Registration of H&E-based tissue annotations to the MALDI-IMS coordinate frame uncovered pronounced compartmental differences in program abundance (Fig. 2a).

**Figure 2:**
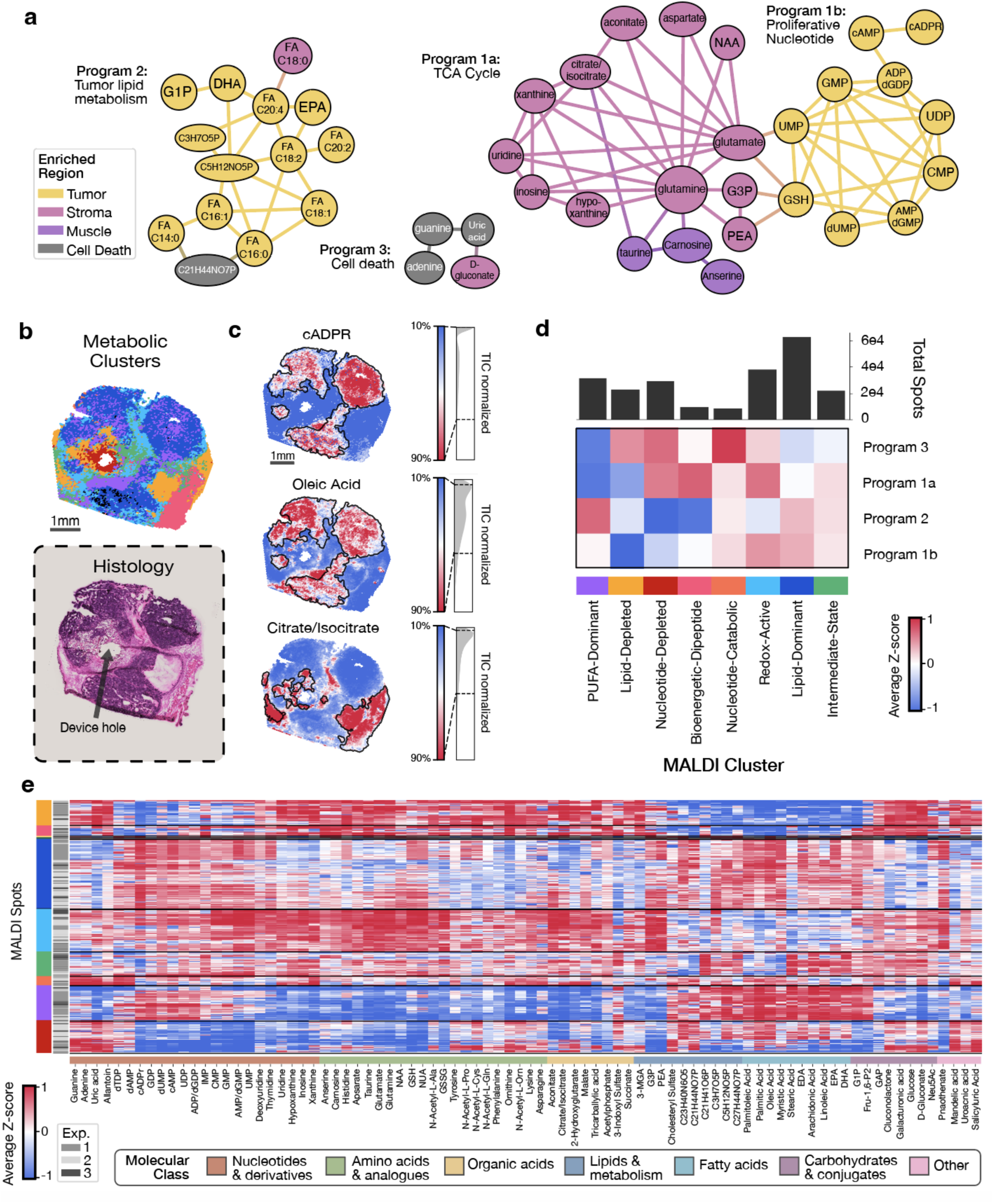
MALDI-IMS-based clustering delineates spatially organized metabolic programs in MMTV-PyMT tumors. (a) Spatial metabolite-correlation network constructed from bivariate Moran’s I (eight nearest neighbors) computed for all metabolite pairs across tumors. Edges denote pairs within the top decile of absolute Moran’s I (retained across all tumor replicates). The network resolves three robust metabolic programs. Nodes are colored by their most enriched pathologist-annotated tissue type (t-test). (b) Comparison of MALDI-derived clusters with histological architecture. Spatial distributions of MALDI clusters are shown across a representative tissue section (top). The central void corresponds to the location of the implanted microdevice used for localized drug release, which was removed prior to sectioning. The matched H&E-stained section is shown for histological reference and comparison of tissue compartment organization (bottom). (c) Spatial distributions of three representative metabolites—cyclic ADP-ribose, oleic acid and citrate/isocitrate—are shown to illustrate their correspondence with MALDI-derived cluster assignments and associated histological features. These metabolites were selected as exemplars of the three major metabolic programs identified above. (d) Mapping MALDI-derived clusters to metabolic programs. Heatmap showing the mean metabolite z-score across all spots for each MALDI cluster–program pair. Spot counts for the eight major MALDI clusters are displayed above. (e) Heat map of z-scored metabolite intensities (rows: MALDI spots; columns: metabolites) highlighting coordinated modules of metabolite abundance that align with spatial segmentation. MALDI spot clustering was performed using Leiden clustering on 84 consensus high-quality metabolite peaks. Each metabolite (n = 84) was z-scored across all laser spots (n = 230,428). Both metabolites and MALDI spots were hierarchically clustered and arranged using optimal leaf ordering. MALDI cluster membership for each spot is indicated by color in the leftmost column (key shown in Fig. 2a), alongside a column denoting the originating mouse experiment. Metabolite relative abundance is shown as z-scores ranging from −1 (blue) to 1 (red). For visualization clarity, MALDI spots were grouped into bins of the 250 most similar spots and averaged. Metabolites on the x-axis are color-coded by molecular class.

The largest component displayed a dual community structure with central hubs glutamine and glutamate. The two poles of this program show clear separation between tumor and stroma: the tumor pole reflects a proliferative metabolic program with features consistent with nucleotide biosynthesis/turnover, signaling intermediates, and antioxidant defenses; the stromal pole suggests catabolic/tissue-remodeling, enriched for nucleotide degradation products (e.g., inosine, hypoxanthine, xanthine, uridine), TCA intermediates (citrate/isocitrate, aconitate), amino-acid-linked metabolites (aspartate, N-acetylaspartate), and membrane-precursor turnover (glycerol-3-phosphate, PEA). In the adjacent muscle, elevated carnosine, anserine, and taurine recapitulated expected skeletal muscle signatures^39,40^. From this data, we speculate regional differences in glutamine utilization between metabolically active stromal and tumor areas, consistent with metabolic symbiosis between compartments^11,41^.

A second component indexed a lipid metabolic program centered on fatty-acid acquisition and remodeling in tumor areas. This program encompassed saturated and monounsaturated fatty acids, which are compatible with membrane biogenesis and energy storage, as well as polyunsaturated fatty acids, including arachidonic acid (AA), eicosapentaenoic acid (EPA), and docosahexaenoic acid (DHA). The presence of these polyunsaturated lipids suggests capacity for lipid-derived signaling (e.g., prostaglandins, resolvins) with potential consequences for inflammation and immune modulation in the local microenvironment^42–44^.

A third, minor metabolic component was enriched in apoptotic/necrotic regions and characterized by purine degradation products (guanine, adenine, and uric acid) along with an oxidative glucose by-product (D-gluconate), consistent with nucleic acid breakdown and a more oxidizing microenvironment^45^.

To map metabolic programs to their spatial organization, we clustered MALDI spots by metabolic profile (Fig 2b,c). We performed principal component analysis (PCA) on normalized metabolite intensities and batch-corrected the resulting embeddings using Harmony^46^ to align samples in latent space, preserving shared biological structure while reducing inter-mouse technical variation (Extended Fig. 5b,c). We then performed Leiden clustering^47^ to the corrected principal component scores, with silhouette analysis identifying 11 clusters as the optimal solution (Extended Fig. 5d,e). Focusing on the eight most abundant clusters (>1% of all spots), we annotated each based on its molecular class composition and enrichment of canonical metabolic pathways (Fig. 2d,e).

### Spatial metabolic state predicts immune cellular neighborhoods across imaging modalities

We tested whether these metabolic profiles encoded reproducible information across modalities using a cross-modal classifier (Fig. 3a). To enable robust cross-modality classification, we restricted analysis to high-quality regions exhibiting accurate correspondence across all three modalities (MALDI-IMS, CycIF, and H&E) (Methods).

**Figure 3:**
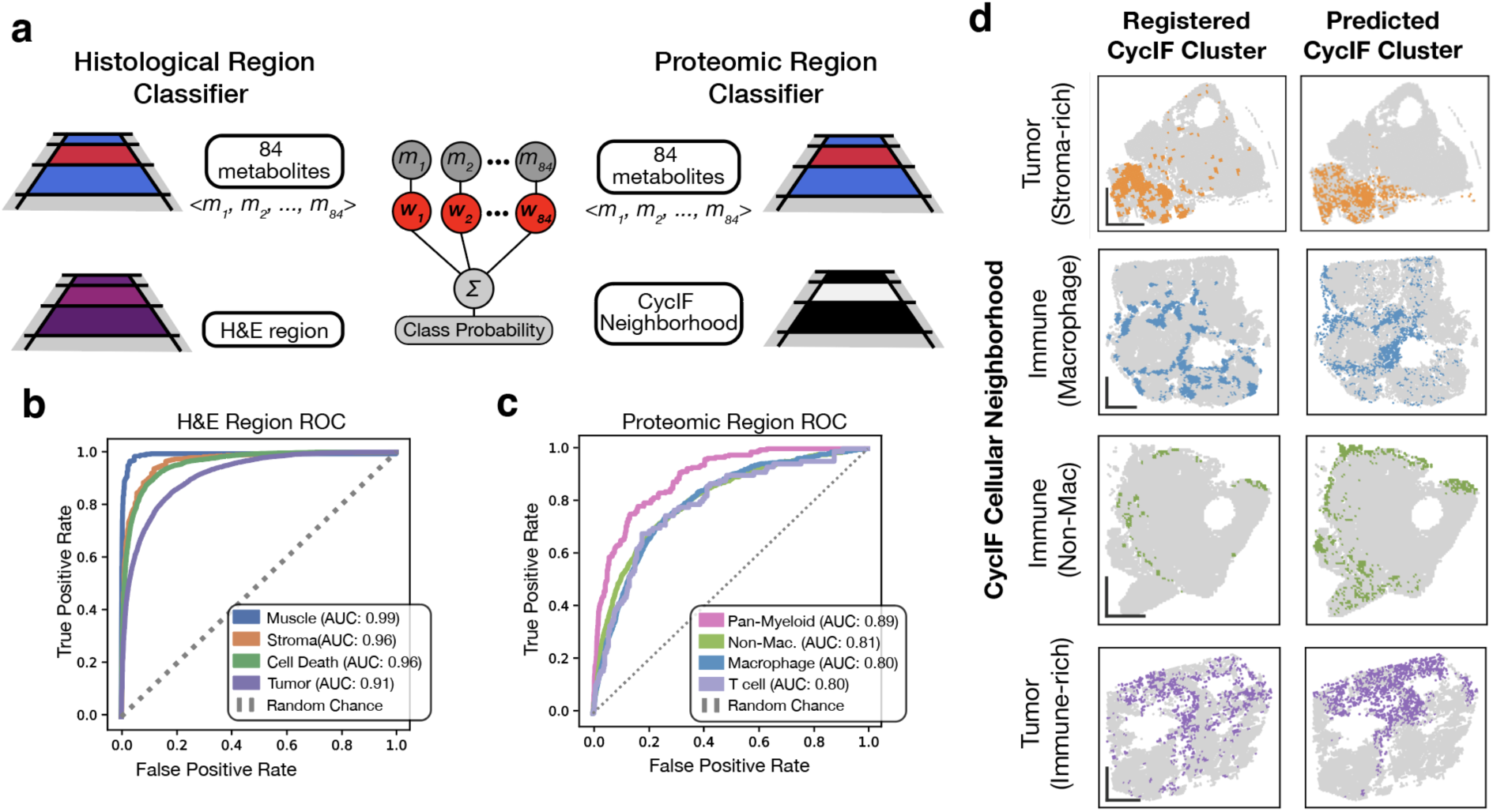
Metabolic profiles predict histology and immune neighborhoods. (a) Schematic illustrating the supervised learning framework. Each MALDI-IMS spot is represented by 84 metabolite abundances, and spot-level labels are derived from registered H&E or CycIF sections. These multimodal registrations generate labeled datasets for model training and for associating classifier coefficients with specific metabolites. (b) ROC curves for logistic regression classifiers predicting H&E-defined histological regions, with performance shown for each class separately. The dashed line indicates random classification performance. (c) ROC curves for logistic regression classifiers predicting CyCIF-derived immune-dominated cellular neighborhoods, shown separately for each class. The dashed line indicates random classification performance. (d) Example of spatial prediction accuracy for a CycIF neighborhood. (Left) Registered dominant cellular neighborhood for the class of interest (“ground truth”). (Right) High-confidence classifier predictions (≥70% probability), demonstrating strong spatial concordance. Scale bars are 1mm.

As a positive control for cross-modality predictive performance, we trained a logistic regression classifier to predict pathologist-annotated histological regions from the H&E section corresponding to the MALDI data (Fig. 3b). Models were trained with class balancing and evaluated using leave-one-section-out cross-validation across 20 tissue sections to prevent overfitting from shared spatial patterns. The model accurately discriminated major histological compartments, with highest performance in muscle (ROC–AUC = 0.98, accuracy = 97%) and lowest in tumor (ROC–AUC = 0.88, accuracy = 81%), yielding a mean AUC of 0.93 across regions. These results demonstrate that metabolic profiles alone robustly capture canonical histological structure across modalities.

Metabolic abundance was sufficient to predict immune cell proteomic phenotypes on a serial section (Fig. 3c,d). We trained a logistic regression classifier to predict the dominant CycIF neighborhood of each MALDI spot (Extended Data Fig. 6a,b). Using similar evaluation, the model most accurately classified myeloid-enriched neighborhoods, with the CycIF immune neighborhoods Pan-myeloid, Macrophage, and Non-macrophage averaging ROC-AUC 0.76 and accuracy 76%. By contrast, the metabolically heterogeneous CycIF Tumor neighborhood was less readily identified (ROC–AUC = 0.66, accuracy = 63%), highlighting the robustness of the identified immune-metabolic coupling (Extended Data Fig. 6c,d).

### Two opposing immunometabolic programs define immune-enriched tumor niches

To resolve structured multimodal variation, we applied sparse nonnegative matrix factorization (sNMF)^48^, which identified a set of interpretable, coordinated relationships between metabolic programs and proteomic features (Methods). We identified five sNMF factors as the optimal decomposition based on cophenetic correlation, dispersion, and H-matrix sparseness metrics (Extended Fig. 7a). A beta parameter of 0.1 provided the best tradeoff between reconstruction error and W sparseness (Extended Fig. 7b). Among the five factors, one primarily captured batch-associated variation (Extended Data Fig. 7c), leaving four major cross-modal latent programs corresponding to tumor, stromal, immune, and cell death states (Fig. 4a, Extended Fig. 7d).

**Figure 4:**
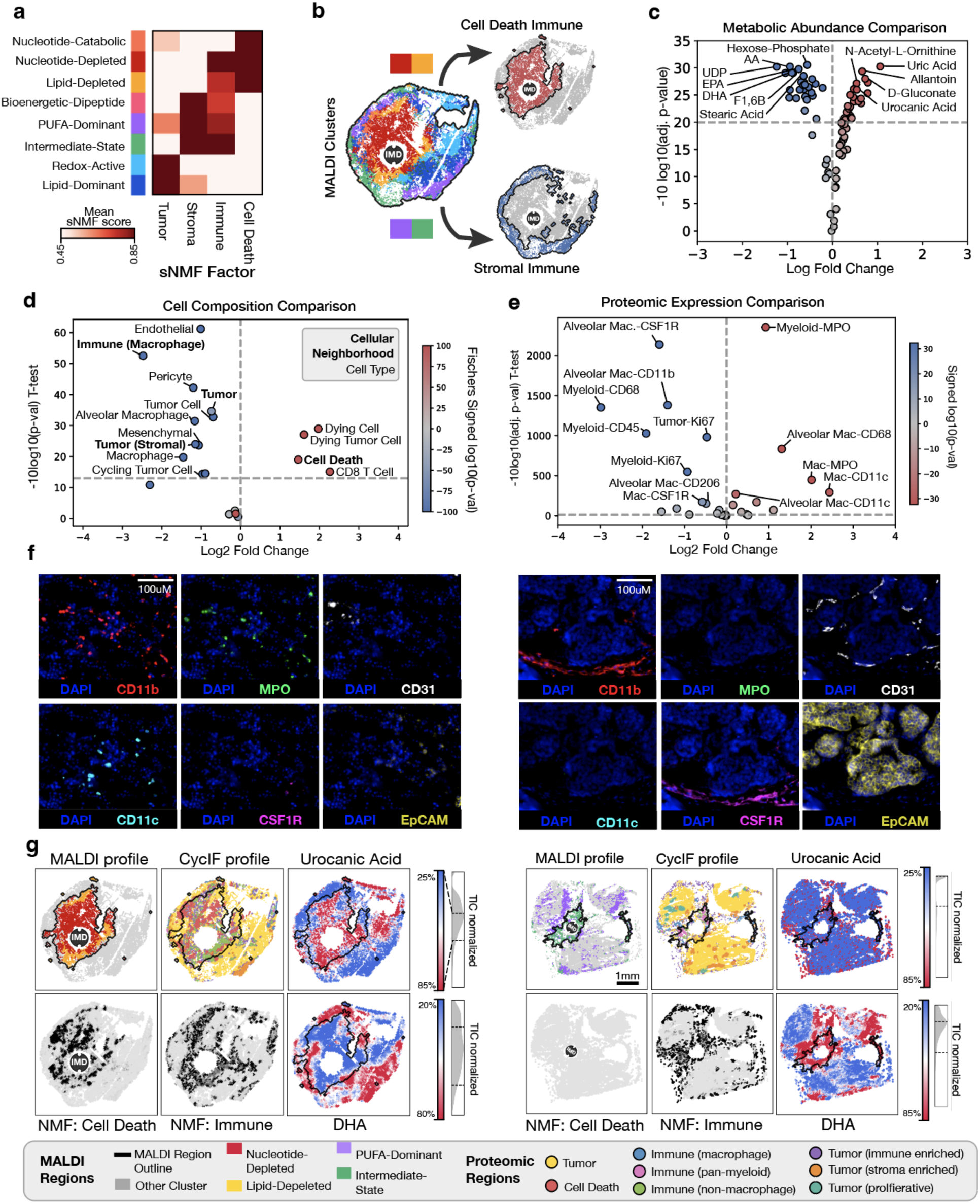
Sparse NMF identifies distinct immune-associated metabolic programs. (a) Mean sparse NMF factor scores across the eight most abundant MALDI-derived clusters. Factor scores were normalized column-wise prior to visualization. Multiple MALDI clusters exhibited immune enrichment, including two clusters additionally associated with strong cell death signatures. (b) Schematic illustrating the splitting of immune-associated MALDI clusters into two major spatial programs: a cell death–associated immune state and a stromal-associated immune state. (c) Differential metabolite abundance between the two immune-associated MALDI programs. Each point represents a metabolite, plotted by log fold change replicates and significance (−10 log of Bonferroni adjusted p-val, LMM). Selected significant metabolites of each program are annotated. Cell death immune metabolites are in red and stroma immune metabolites in blue (Wald test statistic). (d) Differential cellular composition between the two immune-associated programs derived from CyCIF-defined cell types and cellular neighborhoods. Each point represents a cellular identity, plotted by log fold change between regions and statistical significance (−10log10 adjusted P value from pseudobulked t-tests). Immune cell death–associated and stromal-associated regions exhibited distinct enrichments in tumor, macrophage and dying-cell populations. Point color indicates the signed −log10 Fisher’s exact test P value. (e) Differential proteomic cell state markers between the two immune-associated programs. Each point represents a cell type–marker pair, plotted by log fold change between regions and statistical significance (−10log10 adjusted P value from pseudobulked t-tests). Neutrophil- and dendritic cell–associated markers were enriched in immune cell death regions, whereas tumor-associated macrophage markers, including CSF1R and CD206, were enriched in immune stromal regions. Point color indicates the signed significance. (f) CyCIF validation of distinct immune phenotypes associated with the two metabolic programs. Representative fields of view demonstrate differential spatial organization of myeloid, vascular and tumor-associated markers across regions, including CD11b (myeloid cells), MPO (neutrophils), CD31 (vasculature), CD11c (dendritic cells), CSF1R (tumor-associated macrophages) and EpCAM (epithelial tumor cells). Scale bars, 100 μm. (g) Comparison of the spatial mapping of two immune-associated programs. Spatial correspondence between MALDI-derived metabolic programs and CyCIF-defined cellular states is shown for representative immune regions. Representative metabolites associated with each program (urocanic acid and DHA) are displayed as TIC-normalized intensity maps with outlines of the corresponding MALDI clusters overlaid. Spatial distributions of the sparse NMF Immune and Cell Death components are also shown.

We next examined how MALDI-derived clusters aligned with the multimodal sNMF factors and observed clear separation among the eight major clusters (>1% of spots) (Fig. 4b). Two distinct tumor metabolic states emerged, characterized by redox-active and lipid signaling profiles (Extended Data Fig. 8a). In addition, five metabolic clusters were strongly associated with the sNMF immune factor. Among these, the bioenergetic–dipeptide cluster exhibited low overall cellularity and was enriched for metabolites consistent with muscle (e.g., carnosine, anserine, taurine), indicating that its apparent immune enrichment reflects sparse cellularity and extra-muscular myeloid cells. Of the remaining immune-associated programs, two (Nucleotide-depleted and Lipid-depleted) showed elevated cell death factor score (Extended Fig. 8b), whereas two others (PUFA-Dominant and Intermediate-State) aligned with the stromal factor.

To identify key metabolic differences between the two metabolic–immune cluster types, we fit a linear mixed-effects model (LMM) with batch included as a random effect to prevent dominance by any single replicate (Fig. 4c). For additional validation, we computed the pseudobulked log fold changes across tumor replicates and found a near perfect correlation with LMM effect size (Extended Data Fig. 8c). Metabolites enriched within immune-hot cell death regions grouped into several coordinated metabolic programs. These included purine degradation and nucleotide turnover metabolites associated with cellular breakdown (uric acid, guanine, allantoin, adenine, xanthine), oxidative glucose and pentose phosphate pathway metabolites (D-gluconate, gluconolactone, galacturonic acid, glucose), histidine and aromatic amino acid metabolism (histidine, urocanic acid, tyrosine, mandelic acid, 2-hydroxy hippuric acid), TCA cycle and hypoxia-associated intermediates (citrate/isocitrate, malate, succinate, tricarballylic acid, 2-hydroxyglutarate), amino acid and arginine catabolism metabolites (ornithine, N-acetyl-L-ornithine, lysine), and redox buffering metabolites (N-acetyl-cysteine) (Fig. 4c).

Consistent with this redox-driven environment, metabolites most positively correlated with the GSSG/GSH ratio—a classical index of oxidative stress and cellular redox balance^49,50^—included uric acid, mandelic acid, guanine, and adenine. Both MALDI Nucleotide-Depleted and Lipid-Depleted clusters exhibited higher GSSG/GSH than the dominant tumor MALDI cluster (P < 0.01, t-test). Oxidative stress constitutes a dominant axis of variation in the dataset, also captured by PC2 (R² = 0.206 with GSH/GSSG across 230,812 spots), which loads positively on uric acid and negatively on nucleotides (Extended Fig. 8d).

In contrast, immune-rich stromal MALDI PUFA-Dominant and Intermediate-State clusters exhibited broad enrichment of nucleotide pools and nucleotide signaling metabolites, including UDP, ADP/dGDP, UMP, GMP, CMP, AMP/dGMP, GDP, dUMP, IMP, cyclic-ADP-ribose, and cyclic-AMP. These clusters also shared a core metabolic program characterized by abundant lipid and glycolytic intermediates (Extended Data Fig. 8e). Across both states, we observed overlapping enrichment of polyunsaturated fatty acids associated with lipid metabolism (arachidonic acid, DHA, EPA, linoleic acid, eicosadienoic acid), saturated and monounsaturated fatty acids linked to lipogenesis (stearic acid, palmitoleic acid, oleic acid, palmitic acid, myristic acid), and glycolytic intermediates (fructose-1,6-bisphosphate, hexose-phosphate).

Immune composition also differs markedly between the two metabolic cluster types (Fig. 4d). Using pseudobulked aggregation across tissue sections, we found that the MALDI PUFA-Dominant and Intermediate-State clusters are enriched for macrophages (macrophages, alveolar macrophages, and cellular neighborhood *Immune (Macrophage)*) as well as stromal populations (endothelial cells, pericytes, mesenchymal cells, and cellular neighborhood *Stroma Enriched Tumor*). In contrast, the MALDI Nucleotide-Depleted and Lipid-Depleted clusters show increased abundance of tumor cell death (dying cells, dying tumor cells, CD8 T cells, and cellular neighborhood *Cell Death*).

Additionally, immune cells exhibited distinct polarization states across metabolic regions (Fig. 4e,f). Differential expression analysis of single cells stratified by metabolic cluster and cell type revealed that MALDI PUFA-Dominant and Intermediate-State clusters were enriched for TAM-associated markers (CSF1R, CD206) across macrophage populations, whereas MALDI Nucleotide-Depleted and Lipid-Depleted clusters showed increased CD11c and MPO expression across myeloid cells (t-test). Visual overlays of cycIF neighborhoods, sparse NMF components, and metabolic profiles further highlighted these divergent immune phenotypes (Fig. 4g).

### Myeloid polarization state is reflected in reciprocal metabolic programs

To test the robustness of metabolite–protein associations under an orthogonal statistical framework and reduce confounding from cell death–related signals, we performed differential metabolite abundance analysis across CycIF-derived cell types and cell-state markers using linear mixed-effects models with Wald tests and Bonferroni correction (Methods). Network construction from these associations revealed three dominant metabolic–protein poles centered on MPO, CSF1R, and cell death programs (Fig. 5a). Strikingly, CSF1R and MPO showed nearly perfect anti-correlation across metabolites (Pearson r = −0.957; R² = 0.915) (Fig. 5b), defining a metabolic polarization axis that distinguishes TAM-like macrophages from neutrophil-like inflammatory states (Fig. 5c,d).

**Figure 5:**
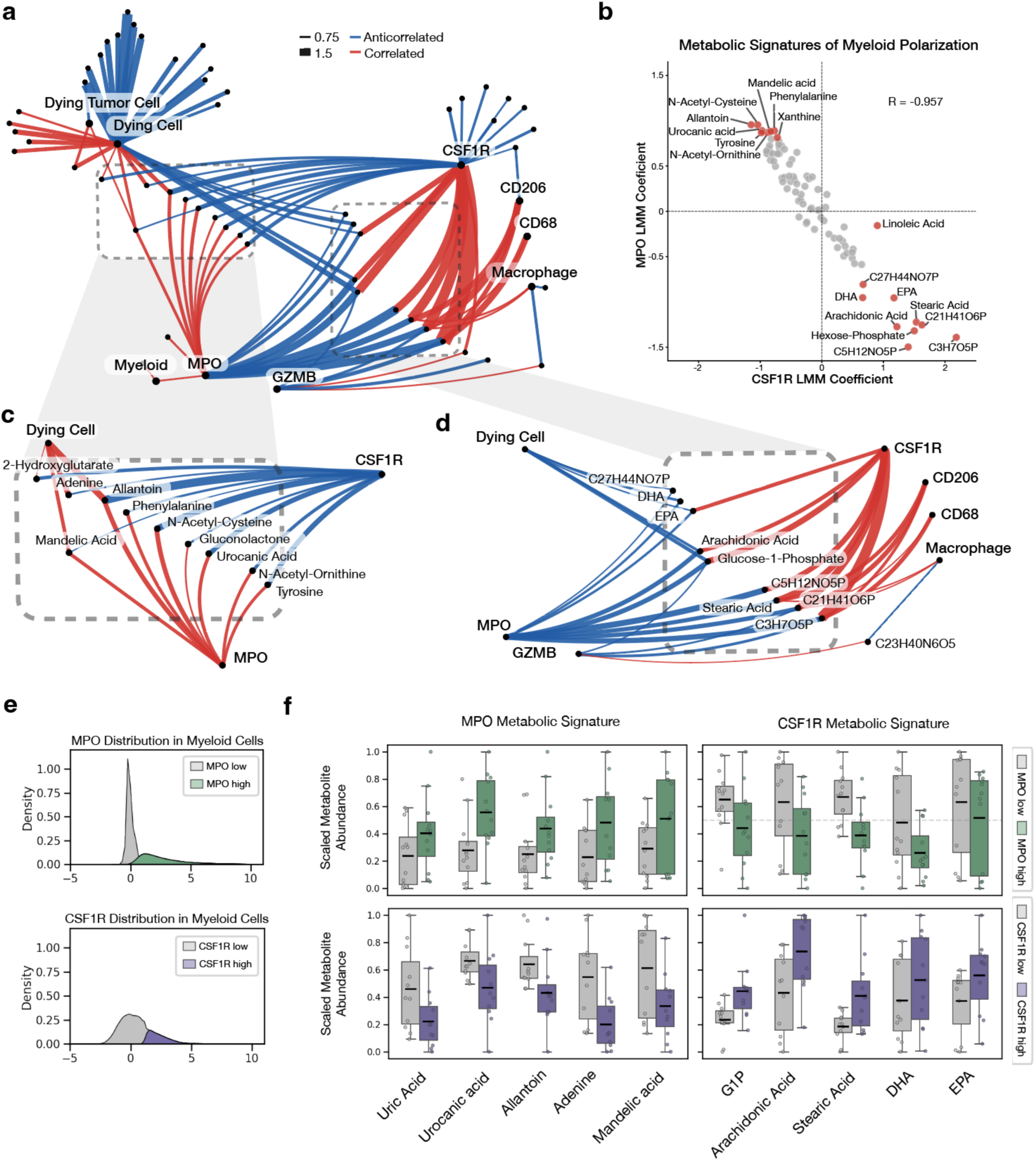
Integrated spatial protein–metabolite networks reveal divergent metabolic programs in MPO- and CSF1R-associated myeloid states. (a) Integrated protein–metabolite correlation network linking spatially resolved metabolites with proteomic markers associated with myeloid polarization and cell death states. Protein marker nodes are labeled. Red edges indicate positive correlations, and blue edges indicate negative correlations, with edge thickness scaled by correlation magnitude. (b) Linear mixed model (LMM) coefficients demonstrating strong anticorrelation between CSF1R- and MPO-associated metabolic signatures across metabolites. Metabolites enriched in MPO-associated regions are highlighted in red and metabolites enriched in CSF1R-associated regions are highlighted in blue. (c) Expanded view of the MPO-associated metabolic network, highlighting enrichment of nucleotide catabolism and oxidative stress-associated metabolites linked to dying cell regions. (d) Expanded view of the CSF1R-associated metabolic network, showing enrichment of fatty acids and lipid metabolism programs associated with macrophage and immune-stromal regions. (e) Distribution of MPO and CSF1R marker intensities across myeloid cells used to define MPO-high/MPO-low and CSF1R-high/CSF1R-low populations. (f) Metabolite profiles stratified by myeloid activation states using GMM-based classification of MPO and CSF1R expression. Differential metabolite abundance was assessed in MPO-high versus MPO-low and CSF1R-high versus CSF1R-low myeloid cells, revealing distinct metabolic programs: MPO-high cells were associated with increased nucleotide catabolism and oxidative stress–related metabolites, whereas CSF1R-high cells showed enrichment of lipid-associated metabolites, including arachidonic acid, DHA, and EPA. Box plots display metabolite intensity distributions across biological replicates, with interquartile range (box boundaries), median (center line), mean (black horizontal line), and individual replicate points overlaid (n=11-15). Each panel shows metabolites previously identified as enriched or depleted in the corresponding cell states. Statistical significance was determined by t-test with FDR correction; all shown metabolites p < 0.05 except for DHA enrichment in CSF1R high cells.

To further assess whether metabolites linked to each marker were consistently associated with marker-positive states, we stratified myeloid cells using Gaussian mixture modeling of marker intensity (two-component fit) and computed section-level mean metabolite abundances within marker-positive and marker-negative populations (Fig. 5e). This approach mitigated inflation of significance from single-cell measurements (Extended Fig. 9a,b). Across this aggregate analysis, metabolites associated with CSF1R- and MPO-defined programs showed coherent directional enrichment patterns consistent with the LMM, with CSF1R-linked metabolites tending to be elevated in CSF1R-positive and reduced in MPO-high states, and vice versa for MPO-linked metabolites (Fig. 5f).

We highlight signature metabolites for each polarized state that were significant in both linear mixed-effects and pseudobulk analyses, exhibiting consistent enrichment in the corresponding myeloid population and reciprocal depletion in the opposing state. CSF1R-associated metabolites included stearic acid, arachidonic acid, EPA, hexose-phosphate, and C5H12NO5P. Whereas MPO-associated metabolites, including N-acetyl-cysteine, N-acetyl-L-ornithine, urocanic acid, allantoin, and tyrosine, showed the reciprocal pattern (Fig. 5f).

### Effective drug treatment reorganizes the local immunometabolic landscape

To evaluate tumor response to treatment, we quantified cell-type abundance as a function of distance from the drug release site (Methods; Extended Fig. 10a,b). Venetoclax induced a strong but inconsistent immune cell recruitment proximal to the device, with immune cells comprising >30% of all cells within 500 μm across three experiments. In contrast, the Panobinostat/Venetoclax (PV) combination—previously shown to have the greatest in vivo efficacy^22,23^—elicited the most pronounced local cell death (Fig. 6a,b). Given the distinct magnitude and phenotypic responses induced by Venetoclax and PV treatment, subsequent analyses focused on these conditions.

**Figure 6:**
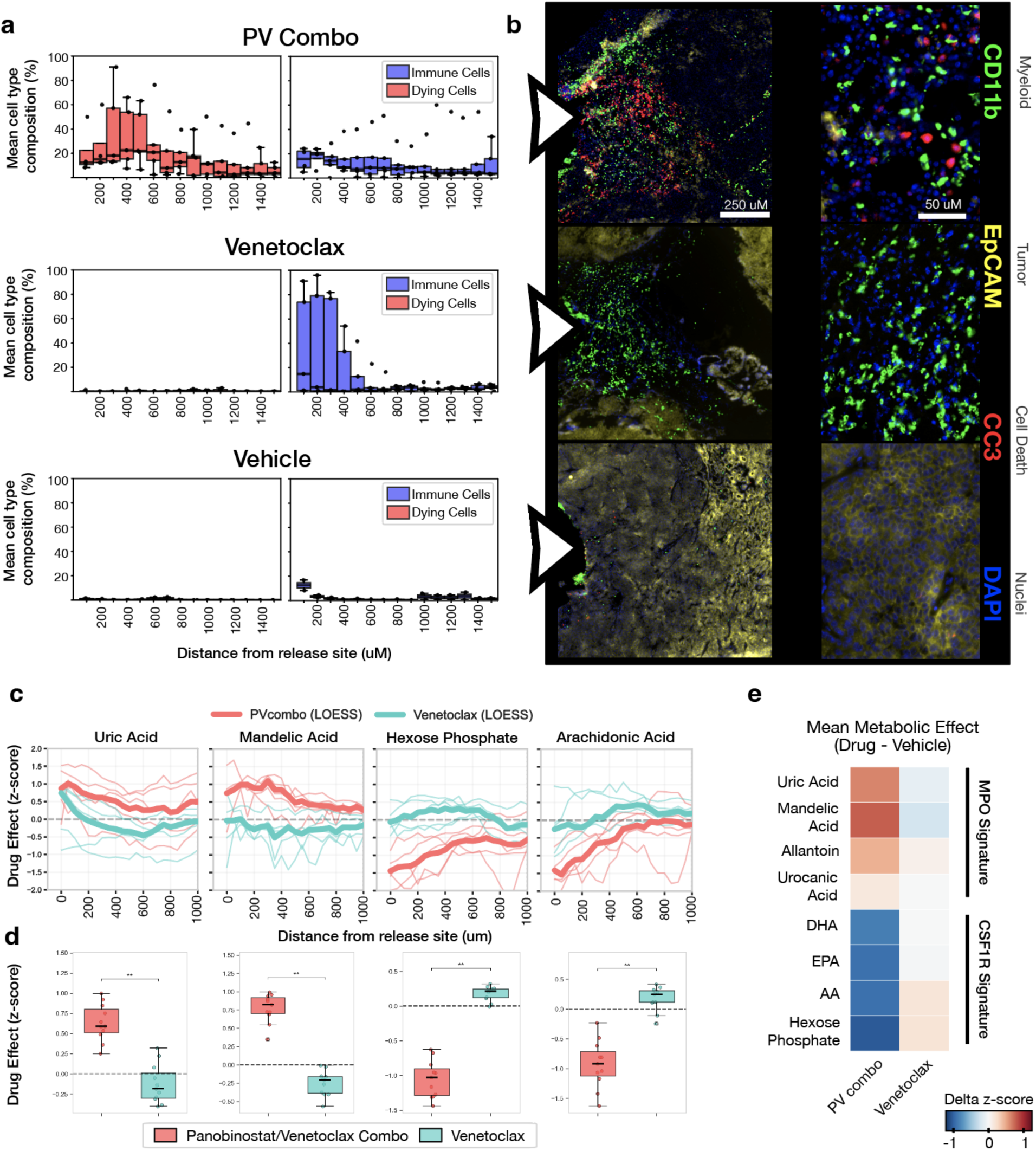
Spatial drug delivery reveals distinct cellular and metabolic responses to panobinostat–venetoclax combination versus venetoclax monotherapy. (a) Recruited/monitored cells at Panobinostat/Venetoclax, Venetoclax, and Vehicle release sites. (b) CycIF images illustrate two prominent induced phenotypes: one elicited by combined panobinostat/venetoclax therapy (a previously validated synergistic pair) and one elicited by venetoclax alone. Myeloid marker CD11b highlights immune infiltration, while CC3 and EpCAM distinguish dying cells from surviving tumor; DAPI labels all nuclei. (c) Metabolite intensities were quantified by MALDI-MSI at increasing distances from implanted drug delivery devices in murine tumors and normalized by metabolite intensity in vehicle-treated regions. Faint lines denote individual tissue sections (n = 3–6 per condition); thick lines show LOESS-smoothed trends (fraction = 0.05). The x-axis spans 0–2000 μm from the drug release, capturing the local diffusion field. (d) Box plots compare metabolite intensities between PVcombo (red) and venetoclax (cyan) within 0–500 μm of drug delivery devices, corresponding to peak drug exposure. Each point represents a section-averaged value (n = 3–6 per condition). Boxes indicate interquartile range with median (center line) and mean (black bar); whiskers extend to 1.5× IQR. Values are expressed as differences from vehicle control to isolate drug-specific effects. Statistical significance was assessed using two-sided t-tests with multiple-hypothesis correction. (e) Heatmap shows mean drug-associated metabolic shifts (drug minus vehicle) for PVcombo and venetoclax across metabolites stratified by myeloid state association. Values represent mean metabolite z-score changes within 0–500 μm of the drug release site.

To determine how immune-associated metabolic states spatially organize under treatment, we performed radial analyses of MALDI cluster abundance as a function of distance from the drug depot (Methods). Tumor cell killing PV combination therapy preferentially enriched for MALDI Nucleotide-Depleted and MALDI Lipid-Depleted clusters, whereas Venetoclax was more strongly associated with MALDI PUFA-Dominant and Intermediate-State regions (Extended Fig. 10c).

To evaluate how these myeloid signature metabolites vary between an effective agent of tumor cell death (PV combo) and an immune recruited less effective treatment (Venetoclax), we tested how individual metabolites vary with proximity to treatment (Fig. 6c,d). Metabolite intensities were binned and averaged by radial distance, normalized to the corresponding vehicle baseline at each distance, and summarized within the 0–500μm treatment zone to enable robust comparison across conditions while minimizing power inflation from large cell count (Fig. 6e). These analyses confirmed spatially restricted treatment-associated changes in metabolite abundance across key metabolites of interest.

## Discussion

This study establishes an *in situ* pharmacology framework that links localized drug exposure to immune-cell states and metabolic organization within intact tumors. By coupling an implantable microdevice with cyclic immunofluorescence (CycIF), MALDI-imaging mass spectrometry (MALDI-IMS), and thin-plate-spline registration, we resolve agent-specific immune phenotypes alongside three reproducible metabolic programs—a tumor-biased proliferative axis, a stromal catabolic/tissue-remodeling axis, and a purine-rich dying cell program—with additional lipid-remodeling features. The venetoclax-proximal increase in myeloid populations, and the PV combination therapy adjacent enrichment for tumor cell death and neutrophil infiltration together indicate that short-range drug exposure is sufficient to reorganize the local immune ecology. Importantly, these cellular shifts occur within distinct chemical neighborhoods defined by MALDI-IMS, positioning immunometabolism as both a readout and a target for rational combinations.

The PUFA-enriched, CSF1R-high, CD206-high, MPO-low macrophage niche identified in our spatial metabolomic analysis is consistent with the lipid-associated macrophage (LAM) state described as an immunosuppressive TAM population in breast cancer. Prior studies have linked LAMs to fatty acid metabolism, TREM2 signaling, immune suppression, and resistance to immune checkpoint blockade^51–53^. Our findings extend this biology in two ways. First, we provide direct spatial metabolomic characterization of a LAM-like niche in intact mammary tumor tissue, demonstrating enrichment of long-chain PUFAs — including arachidonic acid, DHA, and EPA — within CSF1R-high macrophage regions. Second, through pharmacologic perturbation, we show that Venetoclax selectively enriches this PUFA-dominant macrophage state, whereas the PV combination instead promotes an opposing purine-catabolic, MPO-active immunogenic program, suggesting that persistence of a lipid-handling macrophage niche may contribute to Venetoclax resistance.

Fatty acids DHA, EPA, and linoleic acid (LA) were all uniquely enriched in MALDI immune stroma proximal clusters, but not immune enriched MALDI immune cell death clusters. Despite growing interest in supplementing chemotherapies with dietary omega-3 fatty acids, the immunologic functions of DHA and EPA remain incompletely defined in breast cancer. DHA and EPA consistently activate immune regulatory pathways in vitro —including increased IL-10, reduced TLR4 engagement, and dampened NF-κB signaling^43,44,54^. A dietary clinical trial in breast cancer similarly reported DHA-driven IL-10 upregulation^55^, a hallmark of an M2-skewed, immunosuppressive microenvironment and poor prognosis in breast cancer^56,57^. A paired spatial metabolomics and proteomics studies in gastric cancer also reported localized DHA enrichment at tumor–immune boundaries^17^. However, the net impact of these fatty acids on tumor progression remains unclear, with mixed clinical efficacy in breast cancer. Similarly linoleic acid (LA) has been shown to decrease inflammatory cytokine signaling in macrophages through oxylipin signaling in vitro^58^. Our study uniquely links in vivo immunophenotyping in DHA/EPA rich regions with in vitro immune-modulatory effects, providing new insight into how these lipids shape tumor–immune dynamics.

The opposing immunometabolic niche — enriched by the PV combination and characterized by high MPO activity, CD11c induction, and CSF1R downregulation — converges on several established axes of pro-inflammatory myeloid activation. Succinate, elevated alongside malate, citrate/isocitrate, and 2-hydroxyglutarate, is a canonical metabolite of M1-like macrophage activation that stabilizes HIF-1α and promotes inflammatory ROS production^59,60^. Co-enrichment of pentose phosphate pathway intermediates (D-gluconate, gluconolactone) is similarly consistent with elevated NADPH demand during oxidative burst in MPO-active myeloid populations identified by CycIF. Uric acid — the most significantly enriched metabolite in the differential analysis (logFC +1.09) — accumulates during purine catabolism and cell death, where it acts as an immunostimulatory DAMP and inflammasome activator^45,61,62^. Urocanic acid, co-enriched alongside tyrosine, has additionally been linked to reduced MDSC recruitment and enhanced anti-PD-1 responses^63^.

Together, these metabolites define a coherent spatial inflammatory program associated with cell death, dendritic cell infiltration, and M1-like myeloid polarization. Rather than isolated enrichment of individual metabolites, we observe coordinated accumulation of oxidative stress, purine catabolism, and inflammatory signaling pathways within the same tissue niche. This program is pharmacologically distinct from the PUFA-rich, CSF1R-high suppressive niche preferentially enriched by venetoclax monotherapy, suggesting that the balance between these two myeloid metabolic states may influence combination therapy efficacy.

The tumor-biased proliferative program versus the stromal catabolic/remodeling program suggests division of labor across compartments. Our conclusion that tumor–stromal metabolic symbiosis underpins adaptive nutrient exchange and redox homeostasis within the tumor microenvironment is supported by emerging evidence from breast cancer models showing the efficacy of targeting metabolic flux through nucleoside, amino acid, and glucose transport inhibition. Notably, pharmacologic blockade of transporters such as GLUT1 and ENT1/2 has demonstrated potential to disrupt these intercellular nutrient circuits and impair tumor growth in preclinical settings^64^. Further confirmation of this phenomenon requires stable-isotope-resolved metabolomics to distinguish nutrient competition from metabolite-mediated inhibition and to assign flux to tumor versus immune compartments. Glutamine tracing will test whether the stromal–tumor metabolic program 1 reflects compartmentalized glutamine utilization consistent with symbiosis, and whether selective nutrient restriction or metabolite scavenging is sufficient to shift local immune states.

The ability to predict immune neighborhood identity from spatial metabolite abundance alone has not, to our knowledge, been previously demonstrated. Prior integration of mass spectrometry imaging with multiplexed protein imaging has largely been limited to post hoc assignment of metabolic profiles to known cell-type labels^21,65^, and reported limited immune phenotype resolution in a lipid-centric dataset. By contrast, our classifier achieved 76% accuracy for myeloid neighborhoods, respectively, across serial sections without protein information as input. These findings demonstrate that spatial metabolic context is sufficiently structured to encode immune organization. More broadly, these results suggest that metabolite imaging alone may be sufficient to infer local immune architecture directly from clinical tissue sections.

Several constraints temper mechanistic interpretation. First, MALDI-IMS without isotopic labeling is correlative: while the spatial association between metabolites and immune states is robust, it does not, by itself, establish pathway directionality or cell-of-origin. The pixel size and ionization biases inherent to MALDI-IMS mean that signals integrate multiple cell types and extracellular pools; isomer/isobar ambiguity persists for a subset of features, and putative identifications—especially lipids and small polar metabolites-require orthogonal confirmation (targeted LC-MS/MS, ion mobility, or on-tissue MS/MS). Second, although our mixed-effects modeling accounts for within-tumor clustering, multiple wells per tumor are not independent experiments; future designs should emphasize cross-animal replication and prospectively powered comparisons of effect sizes. Third, diffusion kinetics from each reservoir and local tissue microanatomy are potential confounders. At the same time, PEG-only controls mitigate nonspecific effects; more explicit exposure surrogates (e.g., inert tracers, well-to-nucleus distance normalization, and finite-element diffusion modeling) will further constrain dose-response inferences. Fourth, our readouts capture a single exposure window; temporal dynamics (e.g., initiation versus maintenance of the observed immune states) remain to be defined. Finally, our CycIF panel has limited resolution to resolve specific immune populations. While MPO+/CSF1R− cells are consistent with neutrophil infiltration, contributions from other CSF1R-low myeloid populations cannot be excluded without additional lineage markers.

## Online Methods

### Allograft orthotopic MMTV-PyMT

All animal studies were conducted in accordance with protocols approved by the Institutional Animal Care and Use Committee at Brigham and Women’s Hospital. Animals were maintained under specific pathogen-free conditions on a standard 12-hour light/dark cycle with ad libitum access to food and water. The MMTV-PyMT transgenic mouse model on the FVB/N background was obtained from The Jackson Laboratory. Virgin female mice aged 8–24 weeks were used for all experiments.

### IMD Loading

Recommended systemic dose in patients with cancer was derived from the https://rxlist.com web page to June 2017. Systemic doses ranging among 0–1 mg kg^−1^, 1–2 mg kg^−1^, 2–4 mg kg^−1^ and >4 mg kg^−1^ translate to 20%, 25%, 30% and 40% of drug concentration in PEG, respectively, when released from the nanowell. The calibration was determined previously using mass spectrometry measurements^31^. Pure PEG was used in control conditions. Microdevices were implanted for 3 days in MMTV-PyMT with late-stage spontaneously growing tumors in all experiments.

### IMD Retrieval

Implantable microdevices (IMD) were fabricated and utilized as previously reported^22,31^. Briefly, IMDs consisting of cylindrical rods (5.5 mm in length and 750 μm in diameter) were produced from medical-grade polyether ether ketone (PEEK; Solvay) using precision CNC micromachining. Each device contained 18 microreservoirs (200 μm in diameter and 250 μm in depth) arrayed along the surface. Individual reservoirs were filled with therapeutic compounds blended with polyethylene glycol (PEG; MW 1450, Polysciences) as a carrier matrix, while PEG alone served as the control condition.

Devices were implanted into late-stage, spontaneously arising mammary tumors in the MMTV-PyMT model for a duration of 3 days. At the time of implantation, tumors measured approximately 1.2–1.5 cm along the longest axis. Following the treatment period, tumors containing the IMDs were harvested at 3 days post-implantation unless otherwise specified, fixed for 48 hours in either 10% neutral buffered formalin or 4% paraformaldehyde, and subsequently processed for paraffin embedding. Tissue blocks were sectioned using a standard microtome, and 5 μm sections corresponding to each reservoir site were collected for downstream analysis.

### CyCIF Staining and Registration

Before iterative cycles of (1) staining, (2) whole slide scanning and (3) fluorophore bleaching, the slides were subjected to heat-mediated antigen retrieval in citrate buffer (pH 5.5, HK0809K, BioGenex Laboratories, Citra Plus Antigen Retrieval) and then in Tris/EDTA buffer (pH 9.0, S2368, Dako Target Retrieval Solution), all using an electric pressure cooker. Protein blocking was performed for 30 minutes at room temperature with 10% normal goat serum and 1% BSA (BP1600-100) in 1×PBS. (1) Slides were incubated with primary antibody (Supplementary Table 1) for 2 hours at room temperature. All washing steps were performed for 3 × 5 min in 1×PBS while agitating. Slides were mounted with SlowFade Gold antifade mountant with DAPI (S36938) using a Corning Cover Glass (2980-245). (2) Images were acquired using Zeiss Axio Scan.Z1 Digital Slide Scanner (Carl Zeiss Microscopy) at ×20 magnification, after which the coverslips were gently removed in 1×PBS while agitating. (3) Fluorophores were chemically inactivated using 3% H2O2 and 20 mM NaOH in 1×PBS for 30 minutes at room temperature while being continuously illuminated. After protein blocking, samples were subjected to the next round of staining. Whole-slide images were acquired with Zeiss AxioScan.Z1 automated slide scanner (Carl Zeiss Microscopy GmbH). The iteratively digitized images were registered in pixel resolution using piecewise alignment for layers of mosaics (palom) (https://github.com/labsyspharm/palom).

### Cell Segmentation

Cell nuclei segmentation was performed on the CycIF round 1 DAPI stain using the CellProfiler^66^ (v4.26) module IdentifyPrimaryObject with expected nuclei radius 4um to 12um. To improve segmentation, the DAPI image was altered for more accurate segmentation. First, the DAPI image was sharpened with two convolutional filtering steps. A Gaussian filter with kernel size 5uM was subtracted from the image. A Gaussian filter finds the spatially weighted average of kernel. Subtracting the average background of the image normalizes intensity so that there are no bright outlier regions. Next, the Laplacian of the Gaussian (LoG) of the image was subtracted. The Laplacian of the gaussian is a second order derivative operator which identifies rapid change in intensity from one pixel to its surrounding pixels (i.e. edge detection). Subtracting identified edges improves watershed segmentation by distinguishing individual cell boundaries. After image correction, the DAPI images were input to the CellProfiler pipeline.

### Single Cell Marker Quantification

Imaging marker quantification was computed on individual cells using a custom CellProfiler pipeline. Nuclear segmentation is first performed on the DAPI image. Cell boundaries are then approximated by inflating the nucleus by 5uM. The resulting segmentation masks were used to identify individual cells’ nuclear and extra-nuclear mean marker intensity with CellProfiler’s module MeasureObjectIntensity. Cells with nucleus area <5uM^2^ or >250uM^2^ were excluded from further analysis.

### Marker Intensity Gating

For each CycIF experiment, the mean intensity of each protein marker was calculated across all cells. Each gating channel’s intensity distribution was log-transformed and modeled using a three-component Gaussian mixture model (GMM) implemented in scikit-learn^67^. Typically, the first two components captured negative cells with varying levels of tissue autofluorescence, while the third component represented positive cells. Cells with mean intensities above the mean of the third component were classified as positive, with minor manual adjustments made when necessary to align with visually identifiable positive cells. For markers such as EpCAM, which label the majority of imaged cells, positivity was defined as having a mean intensity above the second component. This approach follows established GMM-based gating methods used in previous studies for identifying positive signal.

### Cell Type Assignment

Using the gated marker intensities, cells were assigned to specific cell types based on canonical expression patterns reported in the literature (Supplementary Table 2). Cells identified as autofluorescence-positive were excluded from further analysis. The following cell types were annotated: pericytes, endothelial cells, mesenchymal cells, alveolar macrophages, macrophages, myeloid cells, CD8⁺ T cells, epithelial tumor cells, proliferating tumor cells, dying epithelial cells, and other dying cells.

### Cellular Neighborhood Computation

Cellular neighborhoods were defined based on the local cellular composition within a 50 µm radius of each cell. For each cell, a neighborhood vector was constructed by counting the number of neighboring cells of each annotated cell type. To prevent overrepresentation of highly abundant populations (e.g., tumor cells), neighborhood vectors were normalized by the overall frequency of each cell type within the sample. The resulting vectors were subjected to principal component analysis (PCA) for dimensionality reduction, followed by batch correction across mice using Harmony^46^. Batch-corrected embeddings were clustered using k-means to identify recurrent neighborhood types. Cluster robustness was evaluated by silhouette analysis with bootstrapped confidence intervals.

### Serial Section Registration

TPS registration was performed using the ELD package^32^. Reference points—specific structural landmarks or visually distinct features that could be confidently identified across serial sections and imaging modalities—were manually selected based on conserved tissue structures. Using these reference points, TPS warping generated a mapping that aligned all associated images so that CycIF, H&E, and MALDI-IMS data shared a common x-y coordinate frame. Rigid and affine transformations were also performed using the ELD package, which computes the transformation that minimizes the sum of squared distances between reference points. The same manually selected points used for TPS were applied for these alternative registration methods. In contrast, the naive rigid and affine transformation used only three reference points per image. These three points were chosen from the convex hull of the original set so that they approximately circumscribed the tissue section. The convex hull of each set of points was calculated using the ConvexHull module of the scipy^68^ spatial package.

### Registration Performance Analysis

Hematoxylin and eosin (H&E) staining was performed on serial sections prior to MALDI-IMS and CycIF imaging for nine tissue sections. Each section was annotated by a board-certified histologist to identify multiple region types, including Tumor, Cell Death, Cyst, Stroma, and Muscle. To assess the concordance of histological labels across the two serial sections, three registration methods were tested: thin-plate spline (TPS), rigid transformation, and a naive rigid transformation using three paired points. For each method, the Jaccard index was calculated for all region types. TPS outperformed rigid and closed-form rigid transformations by 19% and 22%, respectively, for within-modality serial-section registration tasks involving histology-based region labeling (paired t-test, P < 0.01). For cross-modality registration tasks aligning CycIF-derived cell death regions with histologically labeled regions, TPS improved performance by 2% and 25% relative to affine and closed-form affine approaches, respectively.

For cross modality comparison, cell death dense regions identified in the H&E images were compared to high intensity CC3 regions in the CycIF data. To identify CC3-positive regions in immunofluorescence images, raw TIFF files were loaded using tifffile and log-transformed to visualize their intensity distributions for manual threshold selection. Thresholding was applied with NumPy, and small holes in the binary mask were removed using morphological operations from scikit-image^69^ (skimage.morphology). The resulting mask was expanded using OpenCV (cv2.dilate) to ensure complete coverage of CC3-positive areas, and final region outlines were extracted using skimage^69^ package function measure.find_contours for downstream spatial analyses.

To assess the concordance of serial section CycIF cell death masks with H&E-derived cell death regions, three registration methods were tested: thin-plate spline (TPS), affine transformation, and a naive affine transformation using three paired landmarks.

### MALDI Imaging Mass Spectrometry

For MALDI Fourier transform ion cyclotron resonance (FT-ICR) measurements, matrix-coated slides were loaded into a slide adaptor (Bruker Daltonics) and analyzed on a solariX XR FT-ICR mass spectrometer equipped with a 9.4 T magnet (Bruker Daltonics). Data was acquired at a resolving power of 120,000 at m/z 500. External calibration was performed before each run using 1 mg mL−1 arginine, yielding mass accuracy within 1 ppm. Endogenous ions at m/z 124.0068 (taurine), 133.0136 (malate) and 145.0611 (glutamate) were used as lock masses during acquisition. The minimum intensity threshold for peak picking was set to 5 × 10^3^. Laser focus was set to small, and the x–y raster width was 50 µm using Smartbeam-II laser optics. Each pixel spectrum was acquired from 200 laser shots at 1,000 Hz. Ions were accumulated using cumulative accumulation of selected ions (CASI) over an m/z range of 70–300 before transfer to the ICR cell for a single scan. Data was collected using FlexImaging software (Bruker Daltonics).

### H&E staining

Meyer and Briggs hematoxylin (Sigma, MHS32) and eosin (Sigma, HT110332) were used for H&E staining. For MALDI-MSI sections, the matrix was removed with ice-cold methanol for 5 min. Slides were then washed in PBS and water, stained with hematoxylin for 15 min, dehydrated in 95% ethanol for 30 s, and counterstained with eosin for 1 min. Sections were further dehydrated in 95% ethanol followed by absolute ethanol, washed in xylene for 2 min, and mounted with Cytoseal 60 mounting medium (Thermo Fisher Scientific, 8310-4). Whole-slide images were acquired on a Hamamatsu NanoZoomer. Anatomical regions of interest were manually annotated by a board-certified pathologist (SA) in QuPath^70^.

### Metabolic Data Preprocessing

All MALDI-IMS data preprocessing is done on a per-slide basis. IsoScope^71^ is used to detect high-quality peaks from the mass spectra of all spots. To label a peak with a metabolite, its mass must be within 2 ppm error of the expected peaks in our list of 186 verified metabolites. Further analysis was completed on h5ad files with the ScanPy package^72^ (v1.10.3). Total ion count (TIC) was used to normalize all spot metabolite intensities by the total sum of all metabolites at that spot. This means that each metabolite will be transformed to a fraction of total metabolite signal detected at that spot. To perform TIC normalization, we used scanpy.pp.normalize_total. An alternative approach, RMS normalization, is performed by scaling each ion intensity by the root of the average arithmetic means of intensities squared at that spot. These methods give comparable downstream results in our dataset. Metabolite intensities were log-transformed after adding 1 as a pseudo count.

During initial data collection, MALDI-IMS sampling boundaries are manually drawn and may inadvertently include non-tissue areas. This results in small observed regions lacking tissue signal, which we refer to as “edge artifacts.” To remove these edge artifacts, MALDI spots are clustered so that groups of artifact pixels can be removed together according to the histology. All data must be z-scored using scanpy.pp.scale so that no one metabolite disproportionately influences the calculation of the principal components and subsequent clustering. Principal component analysis (PCA) is implemented for dimensionality reduction of the metabolite feature space using scanpy.tl.pca. A neighborhood graph (scanpy.pp.neighbors) that describes the distance between different spots’ metabolic profiles is computed using the first 20 principal components. Leiden clusters are computed across a variety of different resolutions using scanpy.tl.leiden. Clusters which correlate to outlier ion intensity sums or obviously tissue-lacking regions are removed. Upon artifact removal, z-scoring, PCA, and Leiden clustering is recomputed.

### MS2 Validation

Mirror plots comparing experimentally measured MS/MS spectra (gray, positive y-axis) with reference spectra adopted from the Human Metabolome Database^73^ (HMDB; red, negative y-axis) for ten metabolites detected on tissue by MALDI-MS in negative ionization mode. Metabolites were first isolated using a 0.1–0.4 m/z quadrupole isolation window to confirm the presence of only the desired [M−H]− precursor ion. Collision-induced dissociation (CID) was performed with collision RF amplitude between 900 and 1,300 Vpp and collision energy between 7 and 12 V (metabolite-specific parameters). Cosine similarity scores between measured and reference spectra are shown for each metabolite. Molecular structures depict predicted (adopted from Metlin) neutral fragments and the corresponding m/z peaks refer to the [M−H]− ions.

### Spatial Correlation Network Computation

To quantify spatial coupling between metabolites, we applied bivariate Moran’s I with eight nearest neighbors to compute spatial correlations for all metabolite pairs. Bivariate Moran’s I analysis was performed using the Pysal^74^ (version 2.4.0). Edges were retained only for metabolite pairs exhibiting mutual spatial correlation (i.e., metabolite x correlated with the spatial lag of y and metabolite y correlated with the spatial lag of x) with correlation stringently defined as falling within the top decile of absolute bivariate Moran’s I (BVI), as determined by permutation-based significance testing to minimize spurious associations. Only pairs uniquely and consistently spatially correlated across all three experiments were included as edges to construct the correlation network (Fig. 2a). This network resolved three robust metabolic programs. It revealed two reproducible anticorrelation relationships centered on Palmitate.

### Metabolic Spot Clustering

For more accurate spot clustering, metabolite z-scores, principal components, and neighborhood graphs are recomputed after all artifact pixels have been removed from the dataset. Harmony^46^ batch correction was used to integrate principal components across multiple mice. An elbow plot was used to determine the optimal number of batch-corrected principal components considered in neighborhood graph construction. Leiden clustering was performed across a range of resolutions (resulting in between 5 and 28 clusters), and silhouette analysis was used to guide selection of an appropriate resolution. For each resolution, average silhouette scores—computed from within-cluster cohesion and between-cluster separation using both cosine and Euclidean distance—were evaluated to identify local maxima and to assess cluster quality. Silhouette plots were additionally examined to ensure reasonable cluster size distributions and to confirm that individual cluster scores were consistent with the experiment-wide average. Fifteen clusters were resolved; four clusters occurring exclusively in a single mouse and representing <1.5% of all spots were deemed artifactual and excluded from subsequent analyses. MALDI-derived clusters were annotated based on their metabolic class and pathway composition.

### Histology Prediction Logistic Regression Classifier

We trained a logistic regression classifier to determine H&E region labels with metabolic input data using the scikit-learn^67^ LogisticRegression implementation (max_iter = 1000). Histology labels for each MALDI-IMS spot were derived from the annotated H&E image. Feature vectors consisted of 84 high-quality metabolite abundances per spot. Spots were retained for modeling only if they met quality criteria across all three modalities—H&E, CycIF, and MALDI-IMS. Data were split into 9 folds by section and trained in a leave-one-section-out manner to avoid inflated test performance from spatial correlates. To ensure balanced representation, each mouse was sampled in equal proportion when training each regional classifier. Reported ROC-AUC and accuracy statistics represent the average across all leave-one-out classifiers.

### Cellular Neighborhood Prediction Logistic Regression Classifier

We performed a classification analysis analogous to the histological region classifier using serial-section CycIF neighborhood data. We implemented logistic regression using scikit-learn, providing 84 metabolite abundances per spatial spot as input features. In this case, label assignment required an additional step, as a single MALDI-IMS spot may correspond to multiple cells in the CycIF image. To obtain a representative neighborhood label for each spot, we mapped all CycIF neighborhoods intersecting a spot and assigned the most frequent neighborhood identity as the spot-level label. Logistic regression models were then trained using the same metabolite feature set, preprocessing criteria, data split, and class-balancing strategy as described for the H&E-based classifier, with performance evaluated using accuracy and ROC-AUC.

### Sparse non-negative matrix factor calculation

We performed sparse nonnegative matrix factorization (sNMF)^48^ with Python’s Nimfa (v1.4.0) package^75^ to determine low rank representations of cross-modal expression patterns. We selected the factorization rank and sparsity regularization parameter for sparse nonnegative matrix factorization (sNMF) through systematic parameter sweeps evaluating both reconstruction fidelity and interpretability. To determine the optimal rank, we performed SNMF across a range of candidate values (k = 2–10) with multiple random initializations and assessed model stability and structure using cophenetic correlation, dispersion, and consensus clarity, alongside sparsity of the basis (W) and coefficient (H) matrices. We identified k = 6 as the highest rank that preserved stable and well-resolved structure across runs while avoiding over-fragmentation, as evidenced by plateauing stability metrics and the emergence of interpretable, moderately sparse factors.

To select the sparsity regularization parameter (β), we conducted a sweep across orders of magnitude (1e^-5^ to 1) and evaluated the tradeoff between reconstruction error (residual sum of squares, RSS) and factor sparsity. Increasing β promoted sparser, more interpretable basis vectors at the cost of higher reconstruction error. We selected β = 1×10^-1^ at the inflection point of this tradeoff, corresponding to the onset of meaningful sparsity in W without a substantial loss in reconstruction fidelity. Together, these parameters yielded a stable and interpretable decomposition that balanced model complexity, robustness, and biological resolution.

### Differential Expression analysis with Linear Mixed Effect Models

To identify metabolites that differed significantly between the two MALDI-defined clusters, we fit a linear mixed-effects model with mouse as a random effect to estimate effect sizes while accounting for inter-animal variability. Wald tests were used to assess significance, with the number of observations conservatively reduced to three effective samples to avoid inflated statistical confidence; Bonferroni correction was applied to control for multiple hypothesis testing. To assess robustness, we correlated mixed-model effect sizes with those derived from a pseudobulked (by-mouse) analysis, ensuring that detected hits were consistent across analytic approaches. In all reported differential analyses, the pseudobulked and mixed-model effect size coefficients had good correlation (R^2^ > 0.9).

### Differential abundance of categorical protein variables

To identify cell types and cellular neighborhoods differentially enriched in immune-rich regions with and without cell death signatures, we performed a comparative analysis. For robust statistical testing accounting for biological replicates, we computed cell-type frequencies at the tissue section level. For each section (and experimental batch) containing >50 cells per condition, we calculated cell-type proportions as the fraction of cells assigned to each Leiden cluster. These per-section frequencies were then used as independent replicates and compared between conditions using Welch’s t-test to account for unequal variances.

Effect sizes were computed as log_2_ fold changes between mean frequencies in each condition, using a small pseudocount (ε = 1×10⁻¹²) to avoid division by zero. Statistical significance was assessed using p-values from the t-tests, with significance defined as p < 0.05 (-log_10_(p) > 1.3). As validation, Fischer’s Exact Test calculation was performed on discrete counts of each categorical variable.

### Differential expression of protein markers

To identify cell type-specific differential expression of protein markers between metabolic microenvironment regions, we performed a cell-level differential expression analysis. Single cells from proteomic imaging data were stratified by their metabolic cluster assignment and cell type identity. We compared two groups of metabolic regions: immune-enriched cell death clusters versus immune-stromal clusters. For each cell type, we analyzed the expression intensity of relevant protein markers, excluding non-specific markers and restricting analysis to biologically relevant marker-cell type combinations (e.g., CD3E, CD4, CD8, and GZMB were only analyzed in CD8+ T cells; Ki67 and pH2AX were restricted to epithelial and proliferating tumor cells). For each marker-cell type pair with sufficient representation (≥5 cells per group), we calculated the log2 fold change between groups and performed Welch’s t-test to assess statistical significance. This approach enabled identification of cell state changes associated with distinct metabolic microenvironments while accounting for cell type heterogeneity. The analysis revealed how specific immune and tumor cell populations modulate their functional marker expression in response to their local metabolic context.

### Distance from dying cell calculation

To quantify the spatial relationship between myeloid cells and regions of cell death, we calculated the distance from each myeloid cell to its nearest dying cell. For each tissue section, cells were classified by type, with myeloid populations defined as cells annotated as ‘Myeloid’, ‘Macrophage’, or ‘Alveolar Macrophage’, and dying cells defined as those annotated as ‘Apoptotic’ or ‘Apoptotic Epithelial’. Spatial coordinates were extracted for all cells in each category. We then employed a k-d tree algorithm (scipy.spatial.cKDTree) to efficiently compute the Euclidean distance from each myeloid cell to its nearest dying cell neighbor. This approach enabled rapid nearest-neighbor queries across the high-dimensional spatial proteomics dataset. The resulting distances were stored for each myeloid cell and stratified by MALDI metabolic Leiden clusters, with particular focus on MALDI clusters Nucleotide-Depleted and Lipid-Depleted, which showed distinct metabolic signatures. Distance distributions were analyzed per Leiden cluster to identify metabolically defined myeloid subpopulations with differential spatial proximity to regions of cell death. Visualizations were calculated in a one-vs-rest fashion. All distances were calculated in pixel units and converted to micrometers using sample-specific pixel-to-micron conversion factors depending on imaging parameters.

### Metabolite correlations with proteomic markers using linear mixed effect modules

To systematically identify associations between cell type abundances, protein marker expression, and metabolite levels across the tumor microenvironment, we employed linear mixed-effects models (LMMs) to account for between replicate variability. For each of the 84 detected metabolites, we performed correlation analyses with both cell type densities and protein marker intensities measured by multiplexed immunofluorescence imaging. The analysis integrated data from 8 tissue sections, comprising over 400,000 individually segmented cells matched to their corresponding metabolic microenvironment. Each cell was assigned metabolite abundance values based on its spatial coordinates using co-registration between proteomic imaging and MALDI-MSI data. We constructed separate LMMs for each metabolite-protein marker pair, treating metabolite abundance as the response variable and protein expression as the fixed effect, while accounting for mouse-specific random effects to control for biological and technical variation. For cell type-metabolite associations, we aggregated cells by type (e.g., alveolar macrophages, CD8+ T cells, epithelial tumor cells) and calculated mean metabolite abundances per cell type per section. The models incorporated patient identity as a random intercept to account for inter-patient heterogeneity. We adjusted p-values for multiple testing using Bonferroni correction. The analysis identified significant correlations (adjusted p < 0.05) between specific immune cell populations and metabolic signatures, revealing coordinated changes in cellular composition and metabolic programming within the tumor microenvironment. Effect sizes were reported as log2 fold changes with adjusted standard errors to account for the limited number of biological replicates and reduce p-value inflation.

### GMM-Based Cell State Classification and Metabolite Pseudobulking

To identify myeloid subpopulations associated with distinct metabolic programs, we applied Gaussian mixture model (GMM) clustering to MPO and CSF1R expression in myeloid-lineage cells (annotated as ‘Myeloid’, ‘Macrophage’, or ‘Alveolar Macrophage’). For each marker, a two-component GMM was fit to scaled expression values, generating low- and high-expression distributions that were used to stratify cells into low, intermediate, and high expression groups. To enrich for transcriptionally distinct phenotypes, only low- and high-expression populations were retained for downstream analyses, whereas intermediate cells were excluded.

For metabolite analysis, metabolite intensities were pseudobulked within each expression group for each tissue section by averaging MALDI signal intensities across cells, requiring a minimum of 200 cells per group per section. Only sections containing both low- and high-expression populations above this threshold were included as paired comparisons. Metabolite abundances were normalized to a 0–1 scale within each metabolite for visualization across features with differing ionization efficiencies. Statistical comparisons between low- and high-expression groups were performed using paired t-tests across biological replicates with Benjamini–Hochberg false discovery rate correction. This analysis confirmed metabolite programs associated with MPO-high neutrophil-like and CSF1R-high macrophage-like myeloid states.

### Metabolite-protein network construction

Metabolite–protein correlation networks were visualized in Gephi^76^ using the ForceAtlas2^77^ layout algorithm to define the initial network structure, followed by manual refinement for visualization clarity. Edges were colored according to correlation directionality (red, positive correlation; blue, negative correlation), and only associations with absolute log fold change > 0.75 were retained. Nodes with one or fewer significant connections were excluded from the final network.

### MPO, CSF1R Signature Identification

Combined signatures were defined from LMM coefficients based on reciprocal extremes: the CSF1R signature comprised metabolites in the top decile of CSF1R coefficients and bottom decile of MPO coefficients, whereas the MPO signature comprised metabolites in the top decile of MPO coefficients and bottom decile of CSF1R coefficients. These stringent criteria define metabolites with the strongest opposing associations across the two myeloid markers.

### Cell Type Distribution Along Drug Gradients

To quantify cellular recruitment in response to locally delivered therapeutics, we performed radial distance analysis from drug injection sites. For each tissue section, drug injection coordinates and orientation were identified and a 120° sector oriented along the direction of drug delivery was defined. Single-cell proteomic data were restricted to cells within this sector using polar coordinate transformation. Euclidean distances from each cell to the injection site were calculated and binned into 2-pixel intervals from 10–50 pixels, with all cells beyond 50 pixels grouped together. Cell type abundances within each distance bin were normalized to the total number of cells per bin. Six cell populations were analyzed: myeloid cells, macrophages, alveolar macrophages, apoptotic cells, apoptotic epithelial cells, and CD8+ T cells. Distance-dependent shifts in cellular composition were visualized using stacked bar plots. For drug-level summaries, data from replicate sections treated with the same drug were aggregated and visualized using boxplots with overlaid replicate measurements. For simplified visualization, immune populations (myeloid, macrophage, alveolar macrophage, CD8+ T cells) and dying cells (apoptotic and apoptotic epithelial cells) were grouped into composite categories.

### Individual Metabolite Gradients Along Drug Delivery Axes

To characterize drug-induced metabolic alterations at the individual metabolite level, MALDI-MSI metabolite intensities were analyzed as a function of distance from drug injection sites. Using the same 120° spatial sectors, metabolite intensities were extracted from processed and z-scored MALDI data. Mean metabolite intensities were calculated within each distance bin and tracked across the drug gradient. Line plots with standard error bands were used to visualize spatial metabolite gradients. To identify treatment-specific metabolic responses, metabolite intensity profiles were compared across drug conditions after normalization (subtraction of Vehicle-treated profiles) to emphasize relative enrichment and depletion patterns. Statistical comparisons between distance bins identified metabolites significantly altered proximal to drug delivery sites, revealing spatially organized metabolic responses to therapeutic perturbation.

### LOESS Smoothing for Metabolite Gradients

To better visualize metabolite concentration trends along drug gradients while reducing noise from individual measurement variability, we applied locally weighted scatterplot smoothing (LOESS)^78^ to the distance-binned metabolite data. The LOESS algorithm (implemented via statsmodels module nonparametric.smoothers_lowess) was applied with a fraction parameter of 0.05, providing local polynomial regression that adapts to the data density at each distance point. This non-parametric smoothing approach preserved important local features in the metabolite gradients while removing high-frequency noise, allowing clearer identification of drug-induced metabolic patterns. The smoothed curves were overlaid on individual section data points to show both the biological variability between replicates and the overall trend of metabolite changes with distance from drug injection sites.

### Metabolic Cluster Enrichment Along Drug Gradients

To assess drug-associated metabolic reprogramming, we analyzed the spatial distribution of MALDI-MSI metabolic Leiden clusters relative to drug injection sites using the same radial sectoring strategy. Distances from each MALDI pixel to the injection site were calculated within 120° sectors and grouped into distance bins. Leiden clusters derived from unsupervised clustering (resolution = 0.50) were quantified within each bin and normalized as percentages of total MALDI spots per bin. Four MALDI clusters (i.e. PUFA-Dominant, Intermediate-State, Lipid-Depleted, Nucleotide-Depleted) associated with the sparse NMF immune factor were tracked across drug gradients. Stacked bar plots and aggregated boxplots across biological replicates were used to visualize treatment-associated metabolic gradients and spatial enrichment patterns.

### UMAP Generation

MALDI-IMS data was represented by a UMAP to show the separation and inter-cluster relationships between the final 11 metabolic clusters. The UMAP was generated with scanpy and included all retained MALDI-IMS spots. The UMAP distances were determined from a nearest-neighbor graph computed from the first 5 batch-corrected principal components of the metabolite data. Labels were assigned according to metabolic cluster.

## Acknowledgements

Davidson, Jonas, Tatarova, and Fraenkel Labs for Discussion

SD - Funding sources. SU2C 3.1416 Convergence, CRUK Grand Challenges,

Jonas - Funding sources: R01CA295683 and R01CA223150

Fraenkel - Connie New

Tatarova - The German Cancer Consortium and Loewe Center Frankfurt Cancer Institute Discovery and Development Program Award

Pathologist – Sanda Alexandrescu

## Author contributions

V.P. and Z.T. contributed equally to this work. Conceptualization: Z.T., S.D., and O.J, Methodology: Z.T., V.P., S.D., L.M. and O.J, Investigation: J.B., A.P., Z.T., J.J., G.G., N.P., Formal Analysis: V.P., Z.T., Visualization: V.P., Resources: Z.T., S.D., E.F., and O.J, Funding acquisition & Supervision: E.F., S.D., Z.T., and O.J Writing, original draft: V.P. and S.D. Writing – review & editing: V.P., Z.T., A.P., G.G., J.J., L.H., N.P., J.B., E.M., E.F., S.D., O.J.

## Competing interests

O.J. is a consultant for Kibur Medical. His interest has been reviewed and approved by MassGeneralBrigham in accordance with their outside interest policy.

## Data availability

### Code availability

https://github.com/vpister/imd_immunometabolism

## Figures, tables, and Extended Data

**Extended Data Figure 1.**
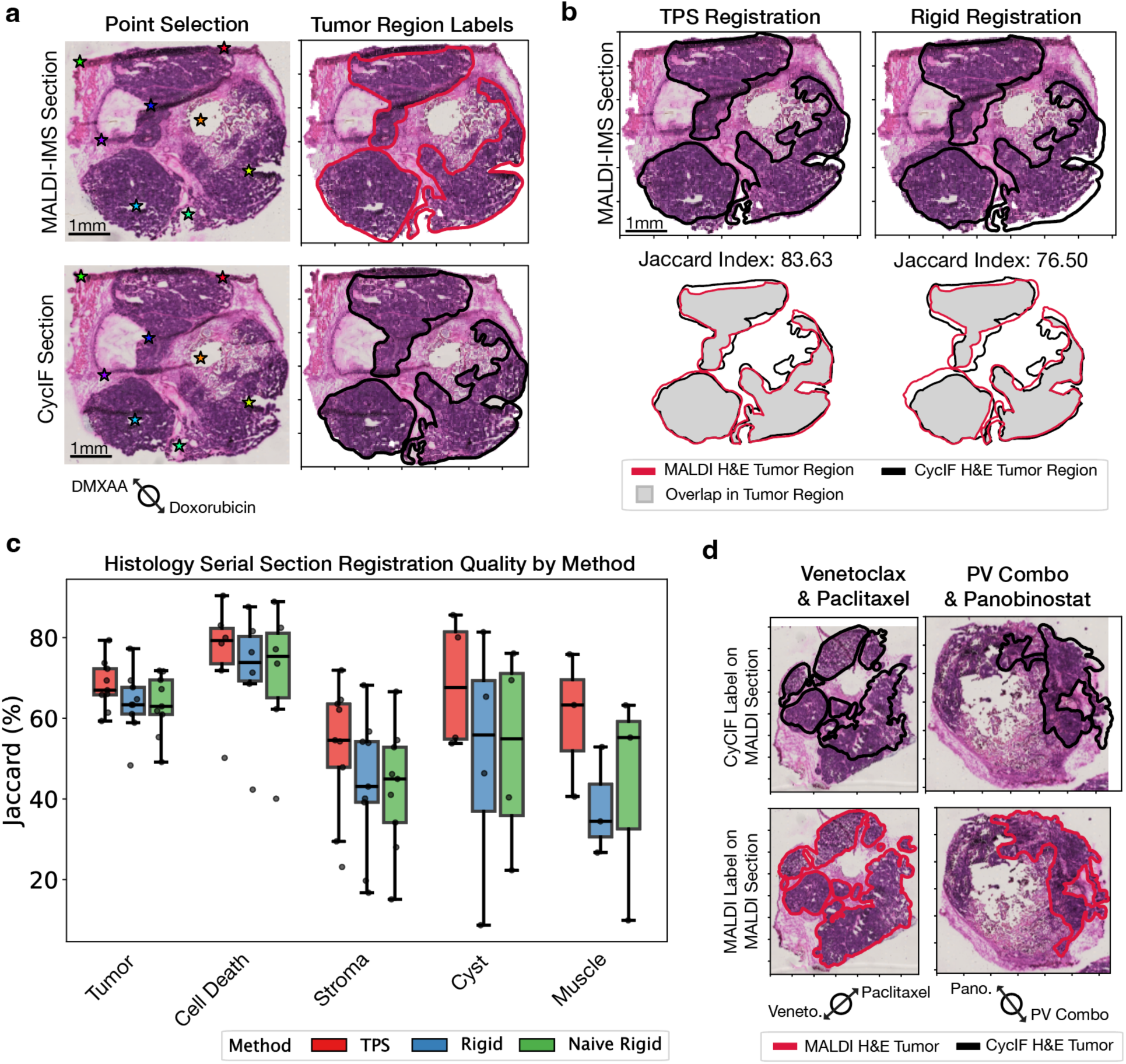
TPS Registration between Serial Sections of H&E. (a) Landmark selection and histological regions from two example serial sections (Doxorubicin, DMXAA treatment) used for MALDI (top) and CycIF (bottom) analysis. Landmarks were manually selected for best fit. Histological annotations were derived from consultation with a pathologist (SA). (b) Comparison of TPS registration and rigid registration using the same set of landmarks. The overlap of same-label regions (Jaccard Index) is used to measure goodness of registration. The region labelled on the CycIF serial section (black) is projected onto the MALDI serial section (top). To visualize overlap, both regions are shown in the same registered coordinate space for both methods of registration (bottom). (c) Aggregated compartmental performance of registration methodologies using Jacard index as a metric. TPS (red) performs the best across different histological compartments. Boxes denote the interquartile range (IQR), with the horizontal line indicating the median. (d) Other examples of registration within the dataset. Registered H&E regions (top) are compared to the ground truth of original labels assigned to the section by pathologist (bottom).

**Extended Figure 2.**
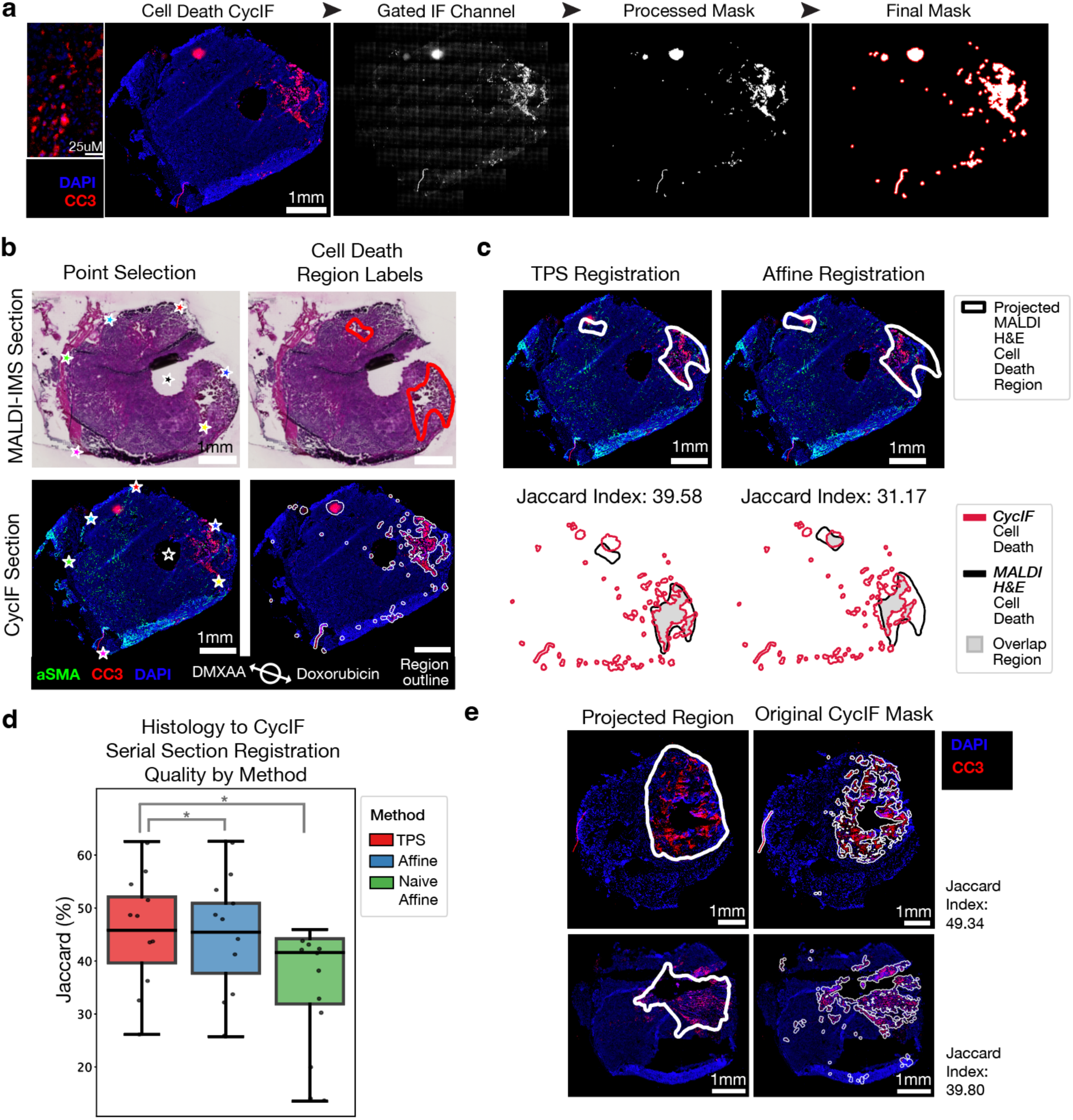
TPS Registration across Serial Sections and Modes of Cell Death Detection. (a) Example of the process of generating masks for cell death channel (CC3) in the CycIF data. (b) Landmark selection and cell death regions from two example serial sections (Doxorubicin, DMXAA treatment) used for MALDI (top) and CycIF (bottom) analysis. Landmarks were manually selected for best fit. Histological annotations of cell death were derived from consultation with a pathologist (SA). CyCIF annotations of cell death were calculated using the algorithm visualized in (a). (c) Comparison of TPS registration and affine registration using the same set of landmarks. The overlap of same-label regions (Jaccard Index) is used to measure goodness of registration. The region labeled on the MALDI serial section is projected onto the CycIF serial section (top). To visualize overlap, both regions are shown in the same registered coordinate space for both methods of registration (bottom). (d) Aggregated performance of registration methodologies using Jacard index as a metric. TPS (red) performs the best. Boxes denote the interquartile range (IQR), with the horizontal line indicating the median. (e) Other examples of registration within the dataset. Registered H&E regions (top) are compared to the ground truth of original labels assigned by algorithmic assignment of cell death regions from the CC3 CycIF channel (bottom).

**Extended Figure 3.**
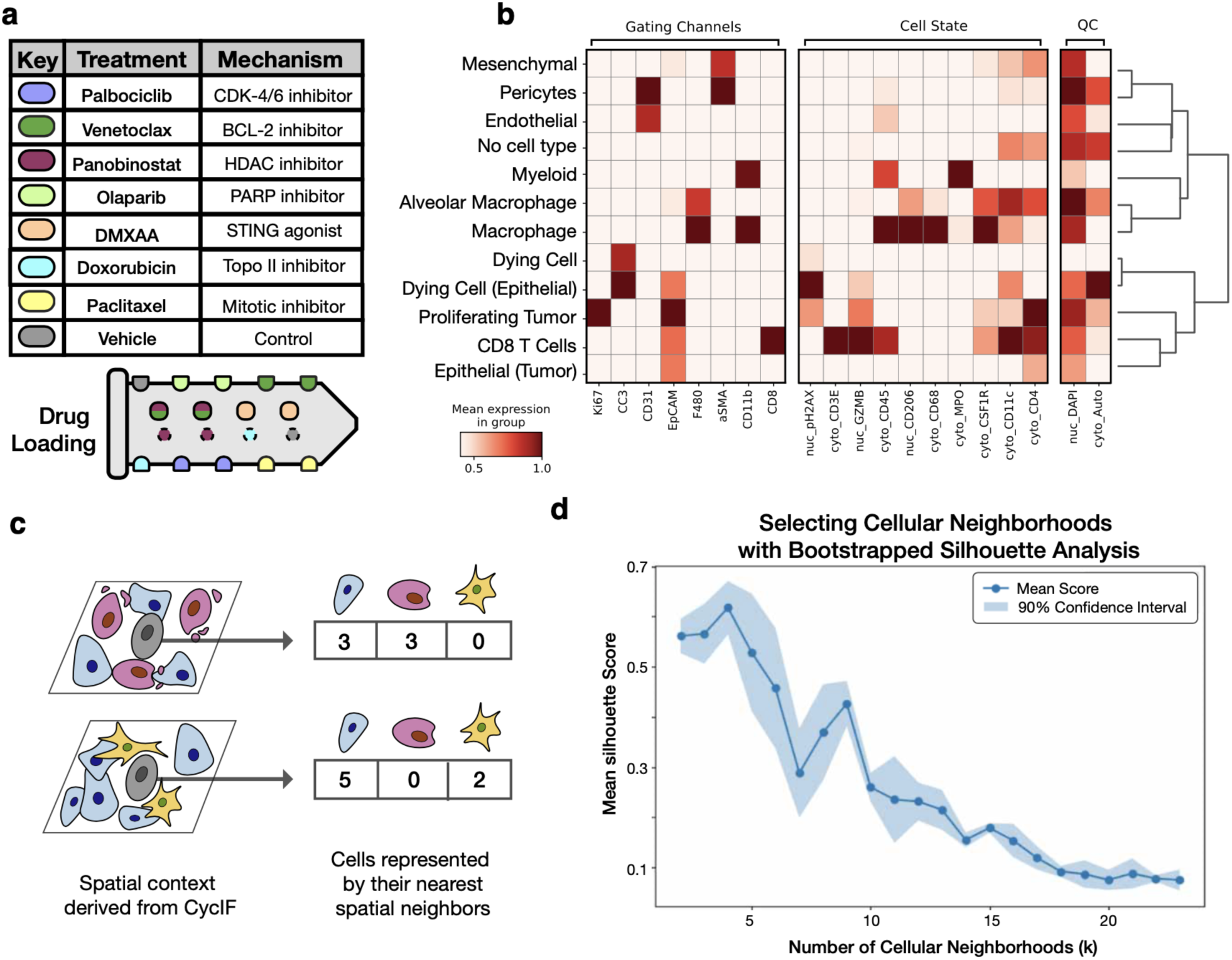
CyCIF-based cell type characterization and clustering. (a) Drug loading diagram and drug mechanisms. (b) Heatmap of mean protein expression across assigned cell types (rows) and CyCIF protein channels (columns). Cell types are hierarchically clustered, with gating channels on the left, followed by cell-state and quality-control channels. Gaussian mixture modeling of individual channels for gating, with cell types annotated according to canonical marker patterns. Expression values are normalized per protein channel, and each cell type shows canonical marker expression. (c) Schematic of cellular neighborhood algorithm. (d) Silhouette plot based on cosine distance, with confidence intervals determined via bootstrapping.

**Extended Figure 4.**
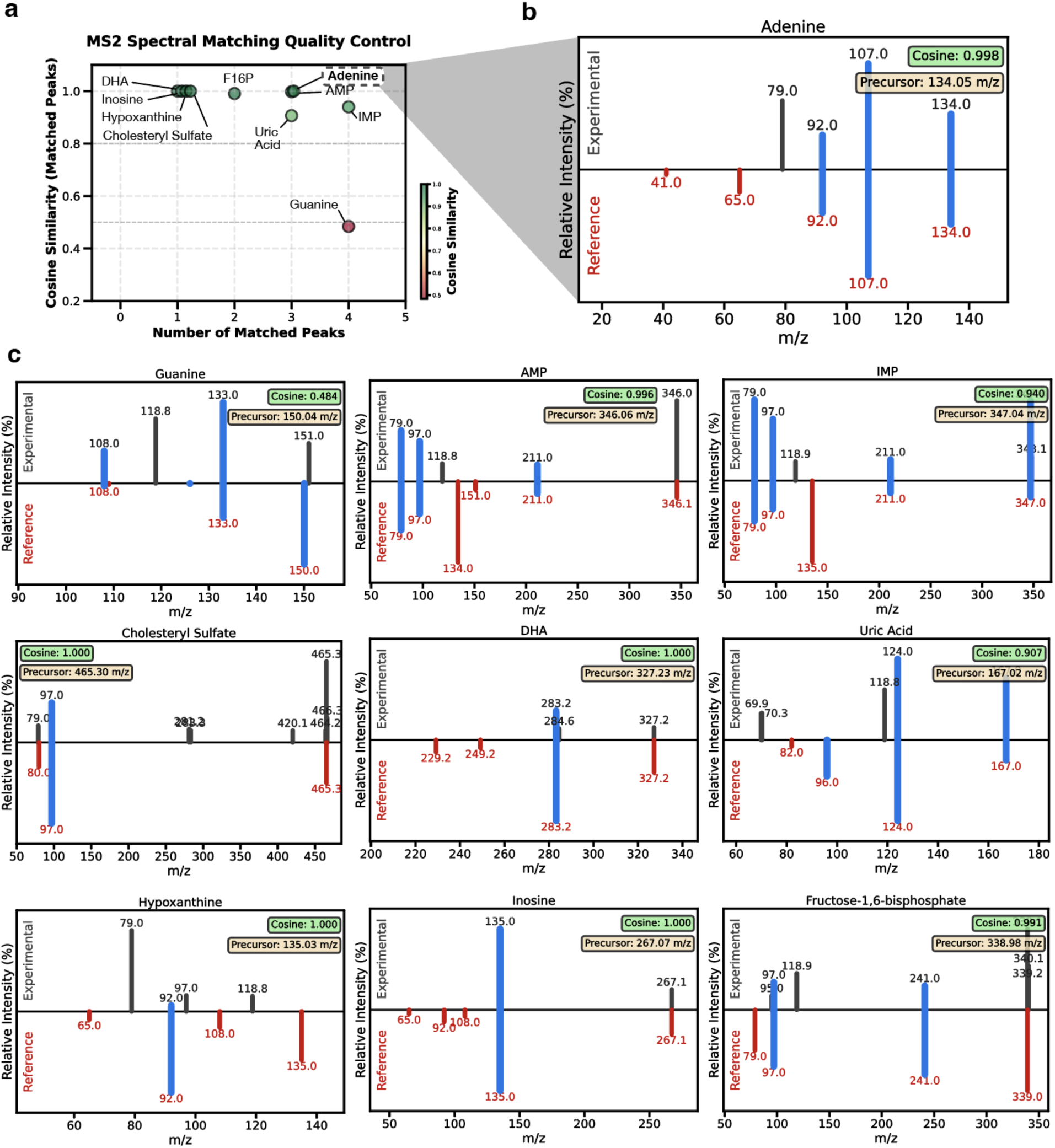
MS/MS Validation of MALDI-IMS peak assignment. (a) Summary of MS/MS validation metrics for annotated metabolites. Cosine similarity of matched peaks versus the number of matched fragment peaks illustrates overall assignment quality, with metabolites in the upper-right quadrant representing the highest-confidence identifications. (b) Representative MS/MS mirror plot for adenine. Reference fragment ions from the library spectrum (red, inverted) are compared with experimentally observed fragment ions (black), with matched peaks highlighted in blue, demonstrating strong concordance between the MALDI-IMS-derived spectrum and the reference standard. (c) MS/MS mirror plots for additional validated metabolite annotations, including guanine, AMP, IMP, cholesteryl sulfate, DHA, uric acid, hypoxanthine, inosine and fructose-1,6-bisphosphate. Reference spectra (red, inverted) are compared with experimentally observed spectra (black), with matched fragment ions highlighted in blue. Most metabolites exhibited strong agreement between reference and experimental fragmentation patterns, whereas hypoxanthine showed comparatively weaker spectral concordance, consistent with a potentially lower-confidence assignment.

**Extended Figure 5.**
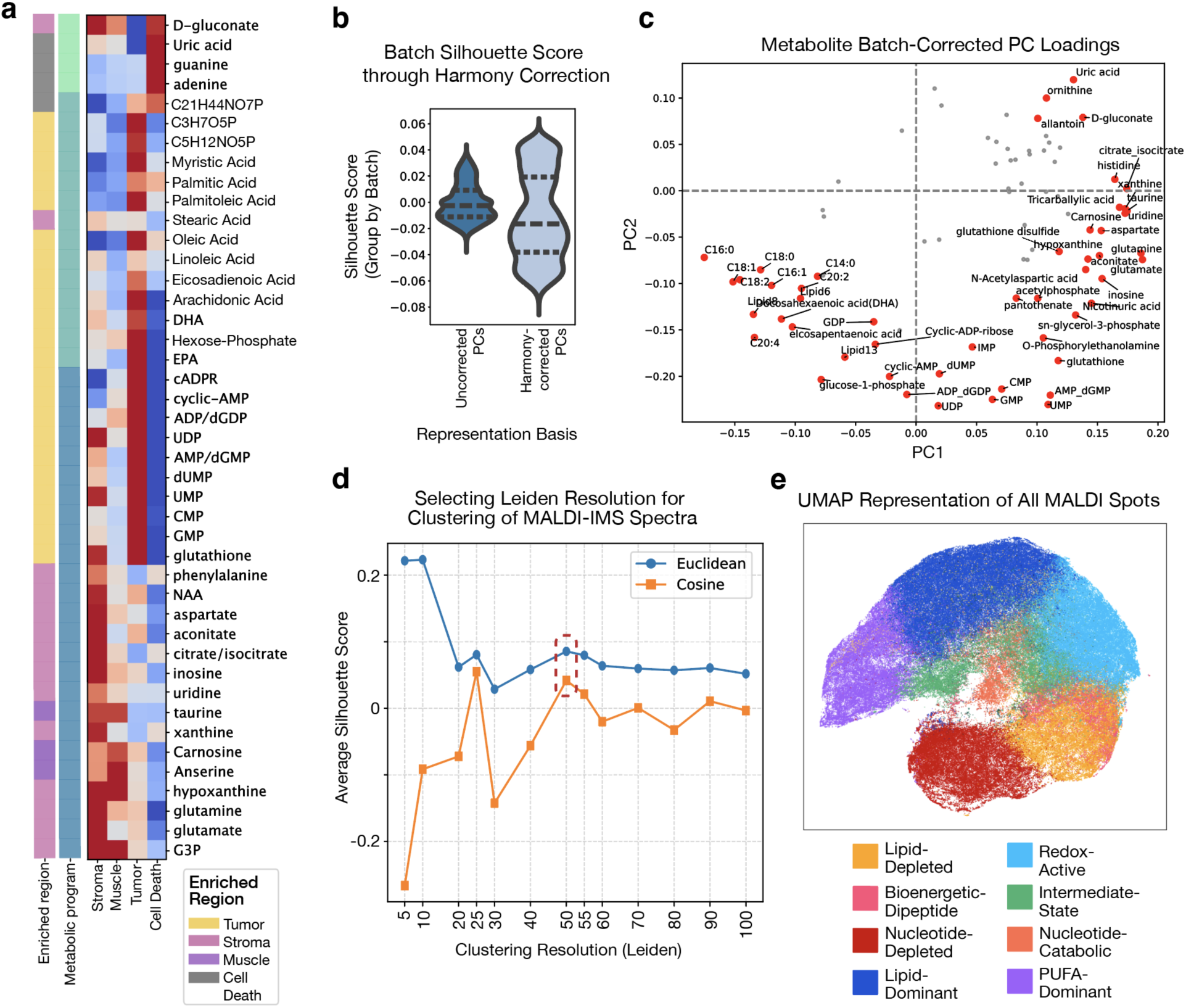
Metabolic clustering segments tissue by robust co-abundant programs. (a) Heatmap of network metabolite components across histological regions. Rows correspond to metabolites, and columns correspond to the histological areas. Colormap values represent t-test statistics. The far-right column indicates the histological region with the most substantial enrichment by t-test, and the rightmost column denotes the associated metabolic program component. (b) Harmony batch correction reduces batch-associated structure in the embedding space. Violin plots show silhouette scores with respect to mouse/batch identity before and after Harmony correction across all spots (n ≈ 230,000). Lower silhouette scores indicate reduced batch-driven separation following correction. Dashed lines denote quartiles. (c) Principal component analysis (PCA) loadings of metabolites. Metabolites are plotted according to their PC1 and PC2 loadings, with those farther from the origin highlighted to indicate greater contributions to the PCs. (d) Silhouette analysis across Leiden resolutions, with scores computed using both Euclidean and cosine distances. Resolution 50 was selected as a local minimum, capturing a suitable number of metabolic clusters to resolve granular immune cell states. (e) UMAP embedding of 230,812 MALDI spots based on Harmony-normalized principal components. Each spot is colored according to one of 11 assigned Leiden clusters. Eight major clusters (>1% of spots are labelled).

**Extended Figure 6.**
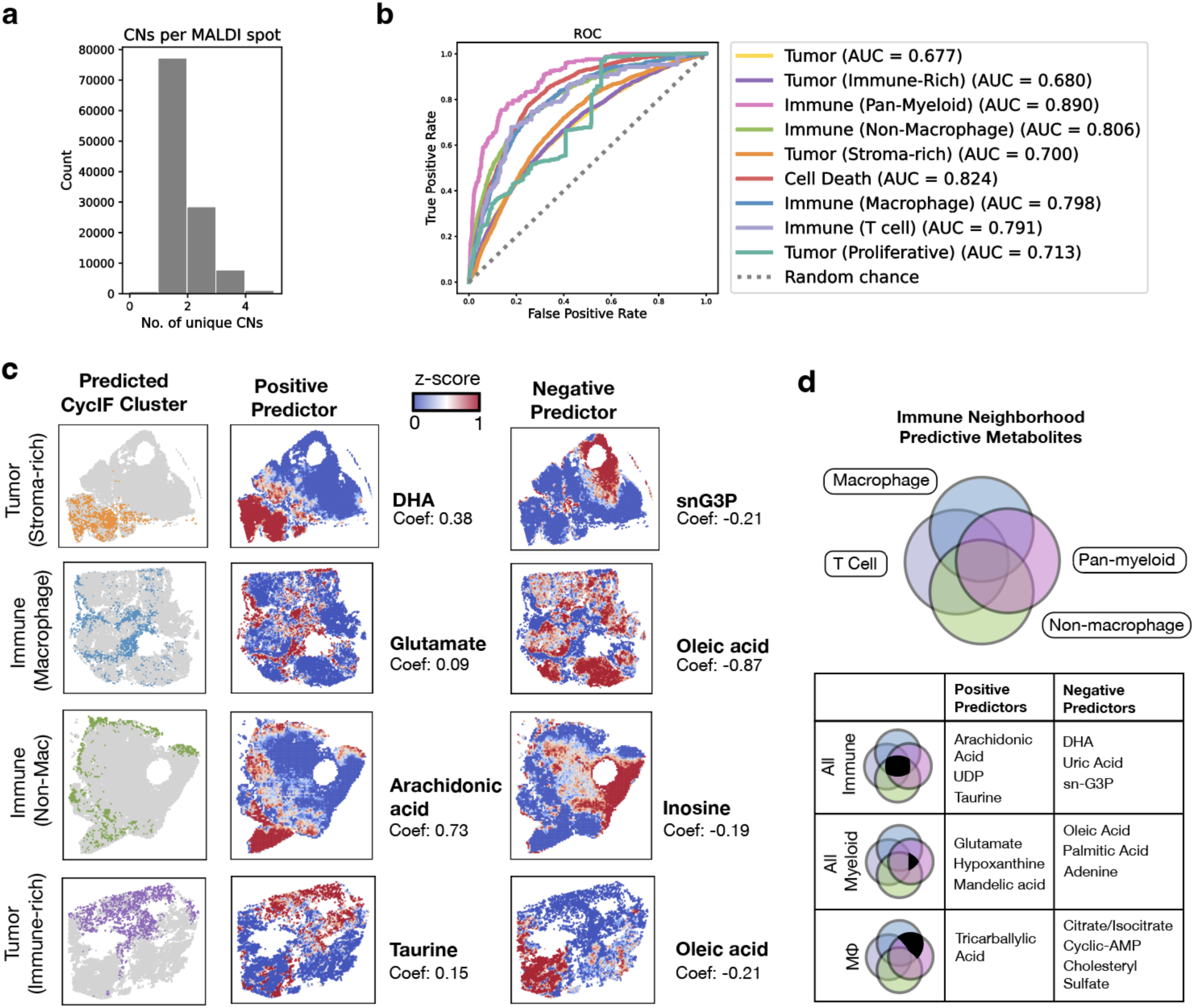
Cellular neighborhood structure and metabolite-based prediction across MALDI-IMS spots. (a) Histogram of cellular neighborhood composition at each MALDI-IMS spot (n=2.3 x 10^5^), showing that the majority of spots are dominated by a single cellular neighborhood. (b) ROC performance of nine cellular neighborhood classifiers across all labels, trained on 84 metabolite features matched to cellular neighborhood identity in a cross-modality, cross-serial-section setting. The dotted line indicates random classification performance. (c) Spatial distributions of example predictive metabolites—one negative predictor (right) and one positive predictor (left)—highlighting the tight correspondence between local metabolite abundance and the mapped immune neighborhood. (d) Venn diagram showing the overlap of predictive metabolites across immune neighborhoods. Representative positive and negative predictors are shown for all immune neighborhoods, for myeloid-enriched neighborhoods, and for macrophage-dominated neighborhoods.

**Extended Figure 7.**
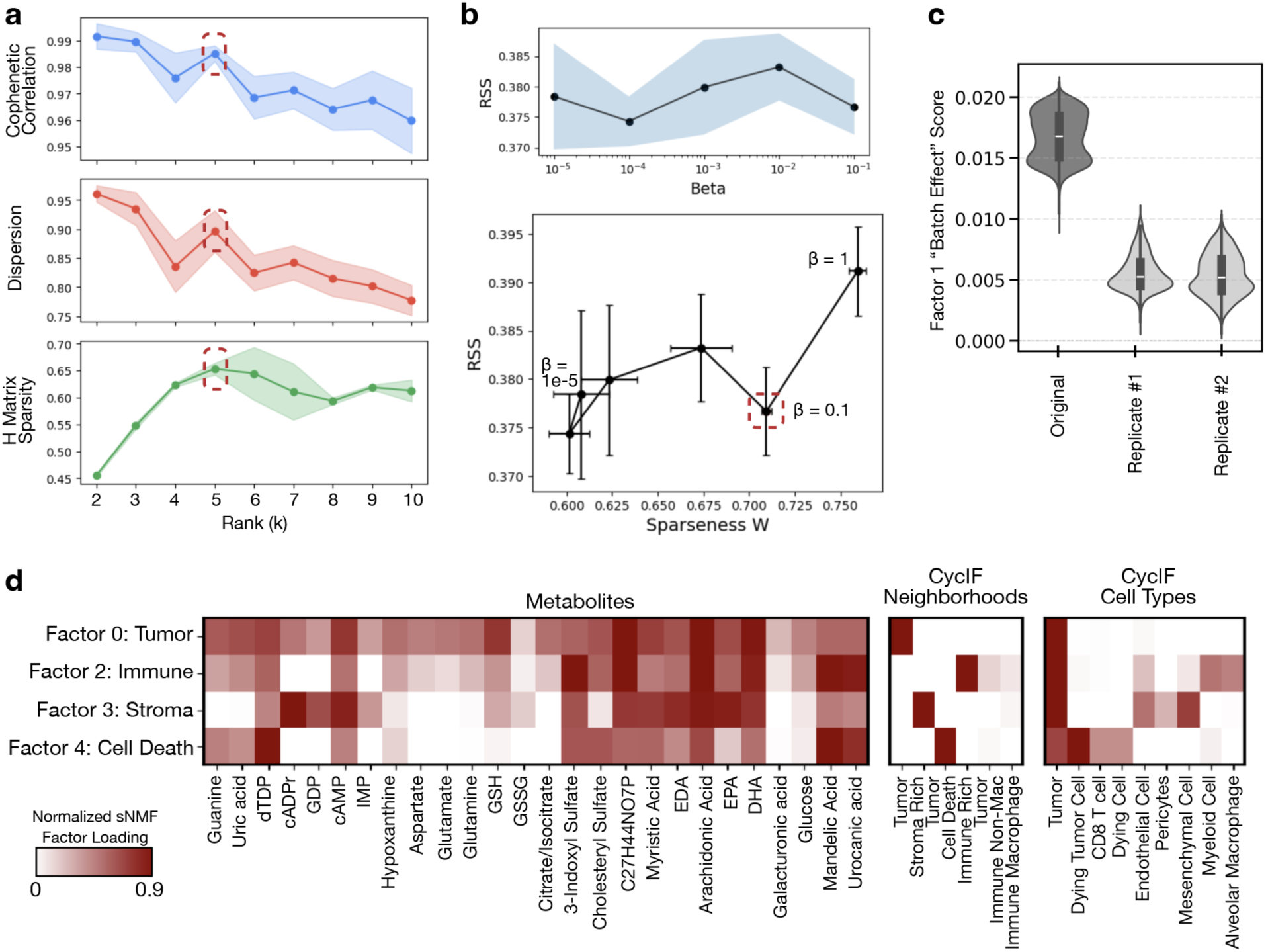
Systematic parameter selection and quality assessment for Sparse Non-negative Matrix Factorization (sNMF) cross-modal analysis. (a) Selection of optimal factorization rank (k). Rank k=5 was selected based on analysis of model stability metrics. The plots show cophenetic correlation, dispersion, and H matrix sparseness across rank options 2–10. All metrics achieved a local minimum or optimum at k=5. Solid lines denote means, and shaded regions represent the 90% confidence intervals from 5 subsamplings (10% of spots each) and 5 different initializations. (b) Selection of the optimal sparsity parameter (β). A β value of 0.1 was selected based on the tradeoff between reconstruction error (RSS, residual sum of squares) and W sparsity. RSS across β values (10^−5^, 10^^−4^, 10^^−3^, 10^^−2^, 10^^−1^ and 1) is shown over five subsamplings and five initializations, with the mean indicated by a solid line and the 90% confidence interval shaded in gray (top). RSS tradeoff with W sparsness is shown over the same β values (bottom). (c) Distribution of Factor 1 scores across mouse replicates. Violin plots represent the distribution of scores, with the mean indicated by a white line. Factor 1 was markedly enriched in the original replicate, which was acquired on a different MALDI-IMS instrument, consistent with a batch-associated effect. (d) Loadings from the final sparse NMF decomposition. Sparse NMF was performed after Frobenius normalization across modalities, and displayed loadings were subsequently normalized per row within each modality. Four non-batch-associated factors were identified, corresponding to tumor, immune, stromal and cell death programs.

**Extended Figure 8.**
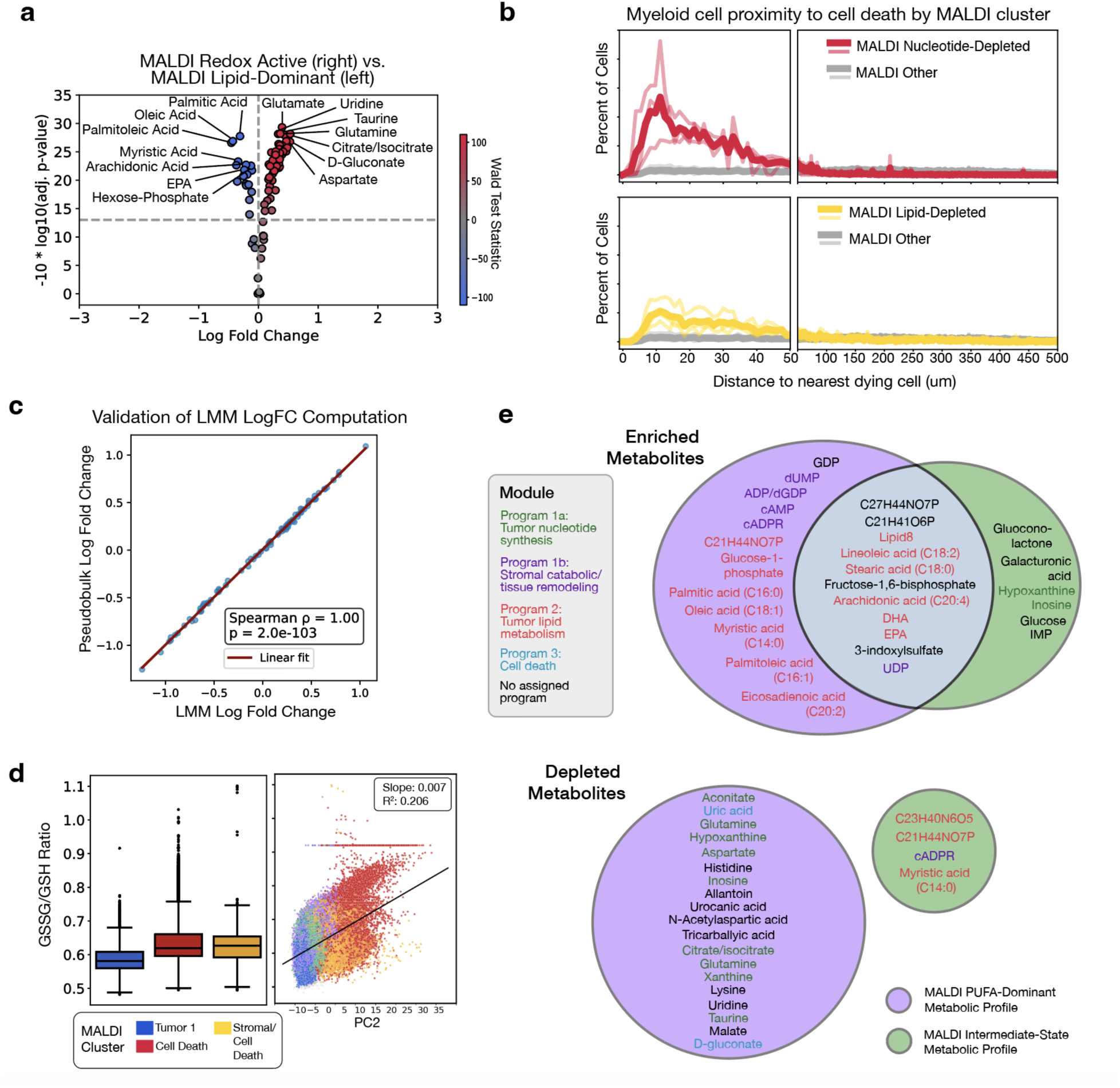
Validation and spatial characterization of MALDI-defined metabolic microenvironments. (a) Differential metabolite abundance between the two MALDI clusters (Redox-Active, Lipid-Dominant) with high sparse NMD tumor factor score. Each point represents a metabolite, plotted by log fold change across replicates and significance (−10 log of Bonferroni adjusted p-val, LMM). Selected significant metabolites of each program are annotated. MALDI Redox-Active enriched metabolites are in red and MALDI Lipid-Dominant metabolites in blue (Wald test statistic). (b) Spatial proximity of myeloid cells to the nearest dying cell across MALDI-defined metabolic profiles. Distances were calculated from each myeloid cell to its nearest apoptotic or apoptotic epithelial cell and stratified by metabolic cluster. Bold lines indicate the mean across replicates and thin lines represent individual mouse replicates (n = 3). MALDI Nucleotide-Depleted regions exhibited the greatest enrichment of myeloid cells proximal to dying cells, followed by MALDI Lipid-Depleted regions. (c) Comparison of differential metabolite abundance between MALDI immune cell death and immune-stromal clusters using linear mixed modeling (LMM) and pseudobulking by mouse. Each point represents a metabolite, with log fold changes independently computed by both methods. Strong concordance between approaches (ρ = 1, P < 0.01) demonstrated robust reproducibility of differential metabolite identification. (d) Redox balance across regions shown as GSSG/GSH ratios (left) and principal component projection (right), illustrating metabolic separation of Cell Death and Stromal/ Cell Death clusters. PC2 from the metabolite PCA correlates strongly with GSSG/GSH (R2 = 0.206 across 230,812 spots). (e) Targeted metabolomics contrasts: both clusters are enriched for fatty acids/lipids (e.g., Linoleic acid, Oleic acid, Arachidonic Acid, DHA, EPA) and tumor-associated intermediates (fructose-1,6-bisphosphate; 3-indoxyl sulfate); MALDI PUFA-Dominant additionally features hexose-phosphate, cyclic-ADP-ribose, and GDP. Depletion patterns diverge: PUFA-Dominant is depleted for uric acid, urocanic acid, allantoin, citrate/isocitrate (mirroring Cell Death enrichments), whereas MALDI Intermediate-State is depleted for C21H44NO7P, cyclic-ADP-ribose, and Myristic acid (tumor-biased species enriched in Mosaic 2).

**Extended Figure 9.**
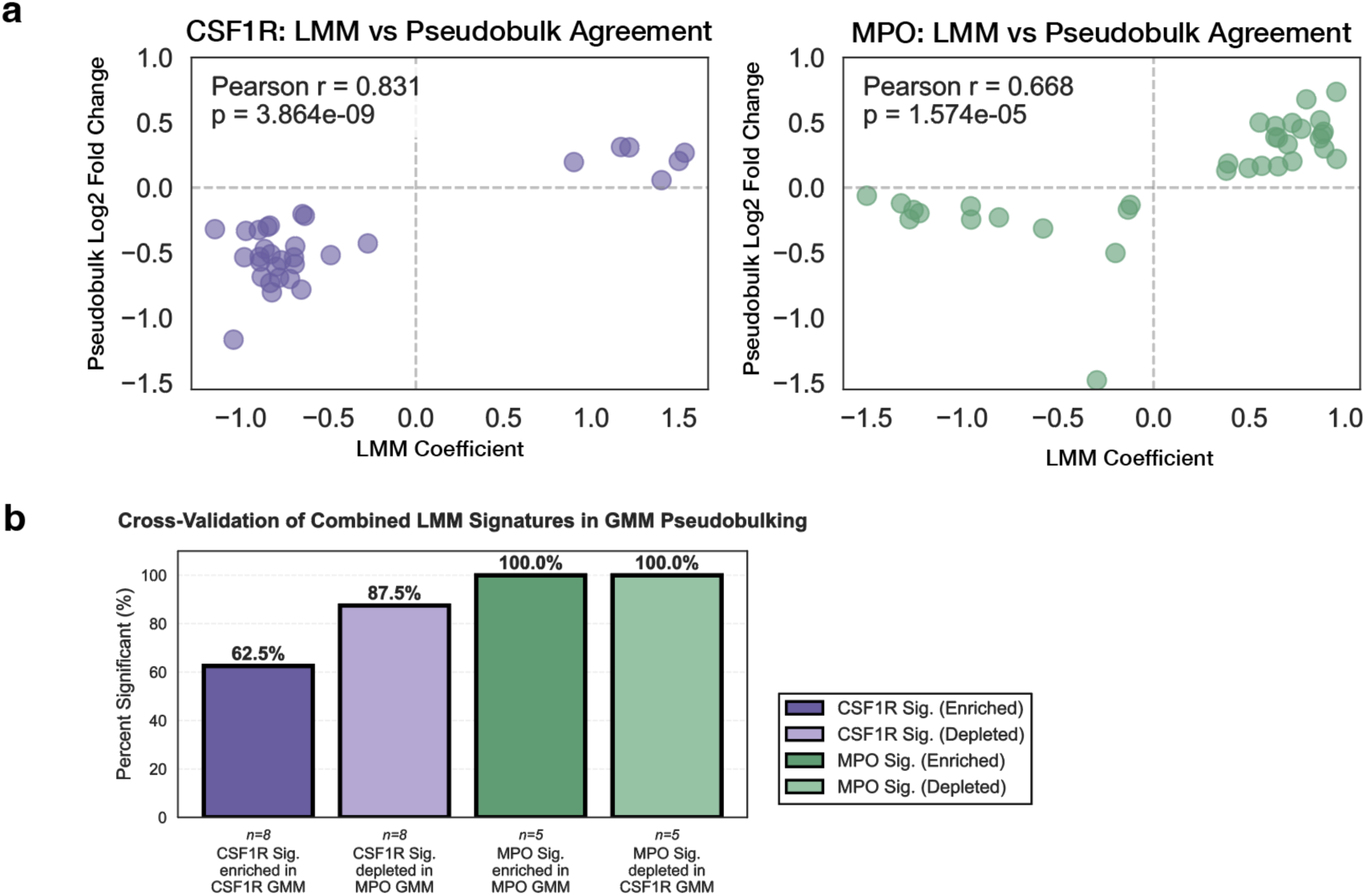
Validating CSF1R-MPO myeloid polarization axis with pseudobulk analysis. (a) Correlations between differential effect size calculations of CSF1R-associated metabolites (left) and MPO-associated metabolites (right). Pseudobulk computation across the tissue section (y-axis) shows high concordance with linear mixed effect coefficient computation (x-axis) in both markers. (b) Cross-validation of combined metabolite signatures between linear mixed-effects model (LMM) and Gaussian mixture model (GMM) pseudobulk approaches. Bar heights indicate the proportion of signature metabolites that were significantly detected (p < 0.05) in GMM pseudobulk analysis with concordant directionality. Solid bars represent expected enrichment (fold change > 1 in marker-high vs marker-low groups), and pale bars represent expected depletion (fold change < 1). Purple bars denote validation of the CSF1R signature in CSF1R-high (enrichment) and MPO-high (depletion) comparisons. Green bars denote validation of the MPO signature in MPO-high (enrichment) and CSF1R-high (depletion) comparisons. n indicates the number of metabolites in each combined signature.

**Extended Figure 10.**
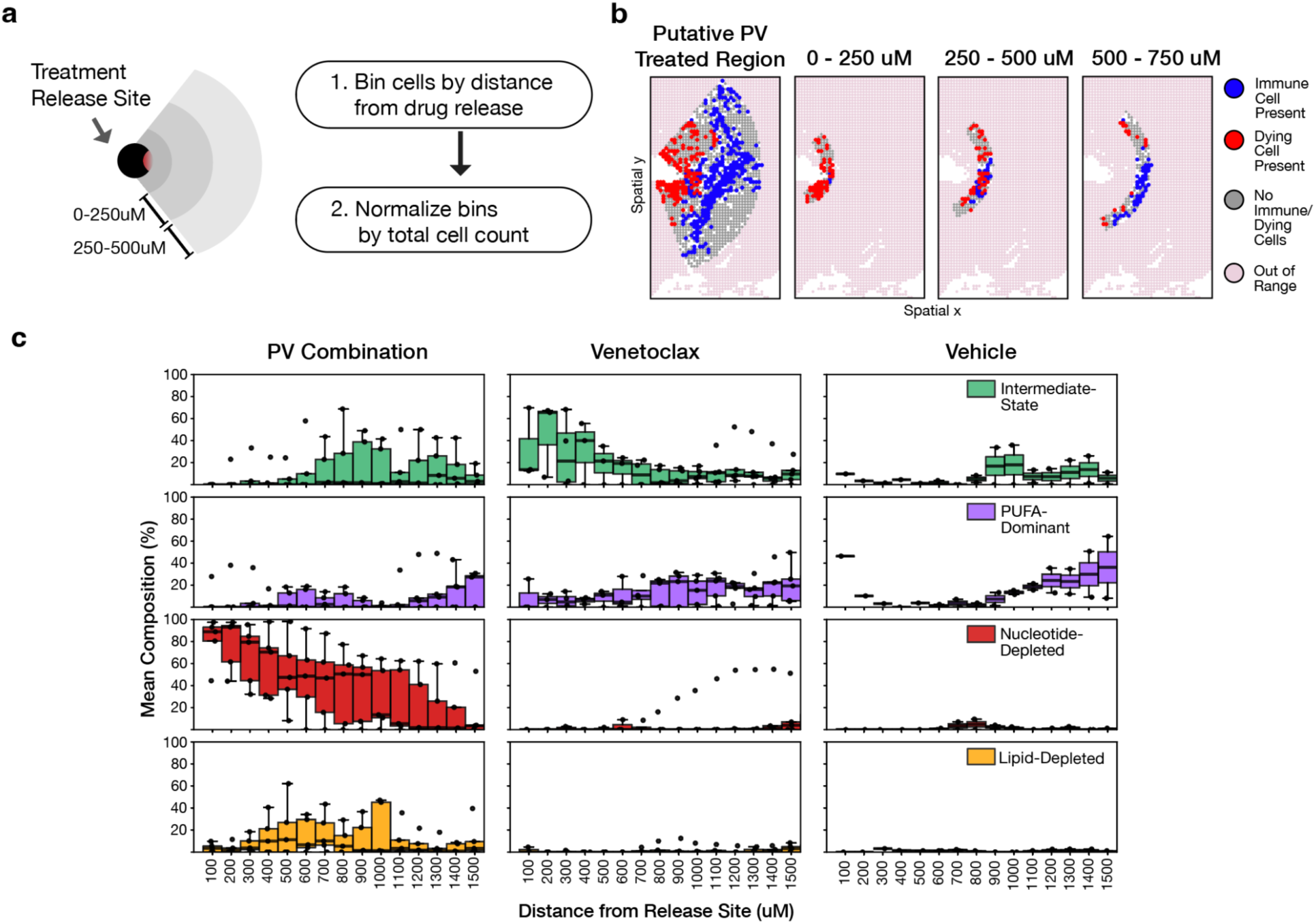
Monitoring Drug Resultant Phenotypes. (a) Schematic of the spatial binning workflow used to quantify drug-associated phenotypic responses as a function of distance from the treatment release site. Cells were grouped into concentric distance bins and normalized by total cell count per bin. (b) Representative spatial map illustrating bin assignment and classification of regions containing immune cells, dying cells, both, or neither across increasing distances from the putative PV-treated region. (c) Distance-dependent MALDI metabolic phenotypes across treatment conditions. Boxplots show the composition of Intermediate-State, PUFA-Dominant, Nucleotide-Depleted, and Lipid-Depleted metabolic programs as a function of distance from the release site. PV combination and Panobinostat treatments preferentially enriched Nucleotide-Depleted regions near treatment sites, whereas Venetoclax was associated with Intermediate-State and PUFA-Dominant metabolic programs. Bold boxes indicate interquartile ranges, center lines indicate medians, whiskers denote 1.5× interquartile range, and points represent outliers.

