## Supplementary Tables for "An in vivo platform to jointly monitor cellular and metabolic responses to chemotherapy"

Supplementary Table 1 | Antibody order, catalog and concentration used in the mouse cyclic immunofluorescence

| **Fluorophore conjugated to primary antibody** | **Vendor** | **Cat No.** | **Panel ID** |
| --- | --- | --- | --- |
| CD11b | Abcam | EPR1344 | Lineage |
| F4/80 | CST | D2S9R | Lineage |
| Ki67 | CST | D3B5 | Lineage |
| CC3 | CST | ASP175 | Lineage |
| EpCAM | Sino | 50591-R002 | Lineage |
| CD8 | eBiosciences | 4SM15 | Lineage |
| CD31 | Abcam | EPR17260 | Lineage |
| αSMA | CST | D4K9N | Lineage |
| CD11c | CST | D1V9Y | Cell state |
| CD206 | CST | 36508S | Cell state |
| CD45 | CST | D3F8Q | Cell state |
| Granzyme B | CST | D6E9W | Cell state |
| CD3 | CST | D4V8L | Cell state |
| pH2AX | CST | 9718T | Cell state |
| MPO | R&D | AF3667 | Cell state |
| CSF1R | Sino | 50059-T24 | Cell state |

Sino, SinoBiological; CST, Cell Signaling Technology

Supplementary Table 2 | Cell type classification decision tree based on canonical markers

| **Cell Type** | **Marker Expression** |
| --- | --- |
| Macrophage | CD11b+ F4/80+ CC3- CD8- CD31- aSMA- |
| Alveolar Macrophage | F4/80+ CD11b- CC3- CD8- CD31- aSMA- |
| F4/80- myeloid | CD11b+ F4/80- CC3- CD8- CD31- aSMA- |
| CD8 T cell | CD8+ CD11b- F4/80- CC3- CD31- aSMA- |
| Pericytes | CD31+ aSMA+ CD8- CD11b- F4/80- CC3- Ki67- |
| Mesenchymal | aSMA+ CD31- CD8- CD11b- F4/80- CC3- Ki67- |
| Endothelial | CD31+ CD8- CD11b- F4/80- CC3- aSMA- Ki67- |
| Dying Cell | CC3+ CD8- CD11b- F4/80- CD31- aSMA- Ki67- |
| Dying Epithelial Cell | EpCAM+ CC3+ CD8- CD11b- F4/80- CD31- aSMA- Ki67- |
| Proliferating Tumor | EpCAM+ Ki67+ CD8- CD11b- F4/80- CC3- CD31- aSMA- |
| Epithelial Tumor | EpCAM+ CD8- CD11b- F4/80- CC3- CD31- aSMA- Ki67- |
